# Recapitulating physiologically relevant oxygen levels and extracellular matrix remodeling in patient-derived tumor-immune tunable models reveal targeting opportunities for immunologically cold high-grade serous tumors

**DOI:** 10.1101/2025.06.23.661098

**Authors:** Simona Plesselova, Hailey Axemaker, Kristin Calar, Oduduabasi Isaiah, Jared Wollman, Somshuvra Bhattacharya, Etienne Z. Gnimpieba, Darci M. Fink, Congzhou Wang, Maria Bell, Pilar de la Puente

## Abstract

High-grade serous tumors are immunologically cold, characterized by limited immune cell infiltration and reduced clinical outcome, primarily due to hypoxia and extensive extracellular matrix remodeling that disrupt tumor-stromal-immune interactions. However, current experimental models fail to fully capture oxygen and matrix microenvironmental features, limiting progress in understanding tumor-immune dynamics and developing effective treatments. Here, we demonstrate that patient-derived tumor-immune tunable models, mimicking physiologically relevant oxygen levels and extracellular matrix remodeling, recapitulate the hypoxia-induced stromal/matrix dysregulation, which causes impaired immune infiltration, and enable dissecting targeted opportunities via TGF-β signaling. The models integrate cancer cells co-cultured with cancer-associated fibroblasts and exposed to immune cells as multi-culture or challenged them to infiltrate into a 3D model bioengineered with autologous plasma from the matching patient or onto decellularized human ovaries. By bioengineering physiologically relevant oxygen levels of hypoxic tumors and physoxic ovaries, we uncovered that intratumoral hypoxia acts as a friend and a foe, causing hypoxia-induced stromal-driven impaired immune infiltration but enhancing the activation and cytotoxicity of CD8+ T cells. We also showed that targeting TGF-β signaling reversed the hypoxia-induced stromal-driven impaired immune infiltration. These human-relevant tunable models may aid the development of targeted therapies to turn immunologically cold tumors into hot ones.

## Introduction

High-grade serous carcinoma (HGSC) is the most common, aggressive, and lethal gynecological malignancy, accounting for 70-80% of ovarian cancer deaths^1^. The five-year survival rate for advanced-stage HGSC patients is less than 30%^2^. The standard-of-care treatment for HGSC typically involves primary cytoreductive surgery followed by platinum/taxane-based chemotherapy^3^. While the initial response to the chemotherapy regimen is usually good, the majority of HGSC patients progress during the initial treatment or recur in less than 6 months after completing treatment^3^. Moreover, ovarian cancer was among the first cancers where the positive correlation between tumor-infiltrating lymphocytes (TILs) and improved clinical outcome was identified in patients with advanced stage^4^. Similarly, the presence of intraepithelial CD3+ and CD8+ TILs is associated with a prognostically favorable survival benefit in HGSC patients^5, 6^. Long-term HGSC survivors present enhanced immune activity with activated memory CD4+ T cells, activated natural killer (NK) cells, and higher densities of CD8+ T cells^7, 8^.

Unfortunately, most HGSC patients are refractory to immunotherapies^9^ and present immunologically cold tumors characterized by immune exclusion from a complex heterogenous immunosuppressive tumor microenvironment (TME) defined by an intricate interplay between the cancer cells, cancer-associated fibroblasts (CAFs), and the extracellular matrix (ECM)^10, 11^. The ECM is the acellular component of the TME that undergoes constant remodeling and provides biochemical and structural support to the cells^12^. CAFs are key players in the dysregulated secretion of ECM proteins through transforming growth factor-β (TGF-β) signaling, which causes a stiff physical barrier for immune cells and drug penetration into the TME^13–15^. Furthermore, the composition and biomechanical cues of the ECM can alter the cellular behavior and drug response of the ovarian cancer cells^16–18^ as well as of the immune cells infiltrated into the TME^19^. Hypoxia, or low oxygen levels, is a key hallmark of tumors that governs CAFs activation and ECM remodeling, interferes in drug resistance, and promotes tumor growth, immune evasion, and metastatic potential^20^. Conversely, hypoxia correlates with an enhanced CD8+ T cell activation, inducing a higher cytotoxic effect of TILs on cancer cells^21^. Although there are neither tumor nor healthy ovarian tissue empirical measurements of the partial pressure of oxygen (pO2), it has been reported that the mean oxygen saturation of human ovarian epithelial cancers is 9.1 percent lower than normal and benign ovaries^22^. Ovarian tumors ranked among the most hypoxic tumor types according to transcriptomic data^23^ and oxygen levels lower than 0.2 kPa were measured in mice ovarian cancer xenografts^24^. In general, hypoxia tends to have median tumor oxygen levels of less than 2 kPa in all tumor types^25^. Physoxic oxygen levels in the healthy human ovary revealed arterial and venous pressures of 14.63 kPa and 11.73 kPa respectively^26^, with follicular fluid dissolved oxygen levels in healthy human follicles ranging from 7.2 kPa to 16.7 kPa^26–28^. Unfortunately, the majority of preclinical cancer models inaccurately compare intratumoral hypoxia to normoxic ambient air 21% O_2;_ however, these levels are hyperoxic with respect to the tissue of origin. In order to better study tumor-stroma-immune cellular interactions and immune infiltration patterns, a more accurate approximation of tissue oxygenation requires hypoxic ovarian tumor levels to be compared to physoxic or physiological oxygen levels of the healthy human ovary.

In this study, we have bioengineered tunable patient-derived models that recapitulate physiologically relevant oxygen levels and ECM remodeling of tumor and healthy ovarian tissue and that are suitable for studying cell-cell, cell-ECM interactions, and immune infiltration patterns. The models rely on HGSC cells co-cultured with CAFs and exposed to peripheral blood mononuclear cells as multi-culture or challenged to infiltrate onto a 3D model bioengineered with autologous plasma from the matching patient or onto decellularized human ovaries. We demonstrated that intratumoral hypoxia drives ECM remodeling through activated CAFs via TGF-ß signaling and alters tumor-immune interactions by collagen signaling pathway with enrichment of immunosuppressive CD4+ T cells and cytotoxic granzyme B+ CD8+ T cells. Our bioengineered tumor models recapitulated hypoxia-induced CAF-driven ECM remodeling and reduced CD8⁺ T cell infiltration, which are hallmarks of immunologically cold tumors. Importantly, TGF-β signaling inhibition with galunisertib reversed immune exclusion and promoted T cell infiltration. Moreover, hypoxia induced higher expression of pro- and anti-inflammatory cytokines involved in tumor progression, which was reversed by targeting TGF-ß signaling. Validation studies in a decellularized ovarian model confirmed hypoxia-induced CAF-driven ECM remodeling significantly impaired immune infiltration, which was rescued by preincubation with galunisertib. Our work provides a paradigm-shifting understanding of intratumoral hypoxia when compared to physiologically relevant physoxic ovarian oxygen levels, allows a mechanistic understanding of hypoxia-induced CAF-driven ECM remodeling and its influence in immune evasion and cellular interactions, and facilities screening of targeting vulnerabilities to reprogram immunologically cold HGSC tumors into hot ones.

## Results

### Single-cell RNA sequencing of patient-derived tunable and physiologically relevant HGSC models identifies intratumoral hypoxia as a driver of ECM remodeling, immune functionality, and tumor-stroma-immune interactions

In order to study the role of physiologically relevant oxygen levels on immune infiltration and immune-cancer-stroma interactions, we bioengineered a set of tunable patient-derived 3D models based on the crosslinking of fibrinogen into fibrin naturally present in peripheral blood plasma^29, 30^ that simulate relevant tissue contexts. Throughout this study, we refer to these as tECMHyp and nECMPhys. tECMHyp stands for tumorous ECM in hypoxia, a 3D scaffold designed to mimic the hypoxic HGSC-TME when incubated in a hypoxia incubator, capturing key features of ECM remodeling in HGSC. In contrast, nECMPhys denotes normal ECM in physoxia, a physiologically relevant 3D model that recapitulates the ECM landscape of the normal ovary under physoxic oxygen conditions achieved when the model is incubated in a standard incubator. The detailed characterization and validation of these models is presented in later sections (see Fig. 1-2 and Extended Data Fig. 1-4). These 3D models were generated to establish a precision-based tumor-immune engineered preclinical model using autologous plasma, peripheral blood mononuclear cells (PBMCs), and primary cells from dissociated tumor biopsies isolated from HGSC patients or HGSC cell lines with plasma and PBMCs from healthy volunteers (Fig. 1a).

**Fig. 1.**
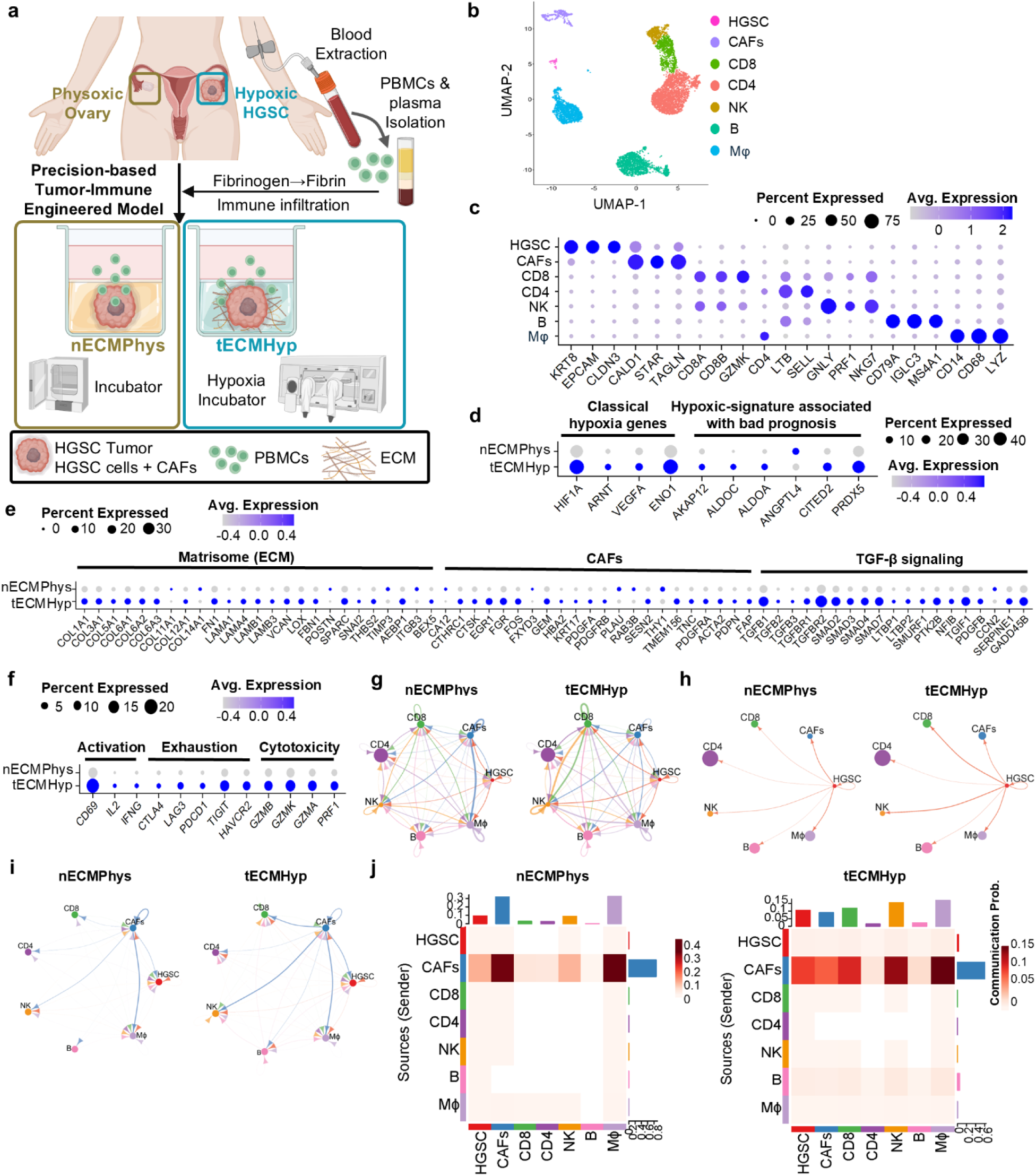
Single-cell RNA sequencing identifies intratumoral hypoxia as a driver of ECM remodeling, immune functionality, and tumor-immune interactions. **a** Schematic of patient-derived 3D model nECMPhys (normal ECM Physoxia) and tECMHyp (tumor ECM Hypoxia) mimicking O_2_ levels in the normal ovary or HGSC, respectively. **b** UMAP plot showing clusters of annotated cell populations. **c** Bubble plot representing the expression of specific marker genes in each detected cell cluster. **d** Bubble plots showing the gene expression of classical hypoxia genes and hypoxic signature correlated with bad HGSC prognosis, **e** matrisome, CAF-associated and TGF-β signaling genes, and **f** T cell functionality genes (activation, exhaustion, cytotoxicity) for nECMPhys vs tECMHyp. **g** Circle plots of interactions (weights/strength) among all cell populations, and **h** HGSC cells with the rest. **i** Circle plots of interactions (weights/strength) among cells in the collagen pathway. **j** Heatmap representing the probability of communication among cells in the collagen pathway.

**Fig. 2.**
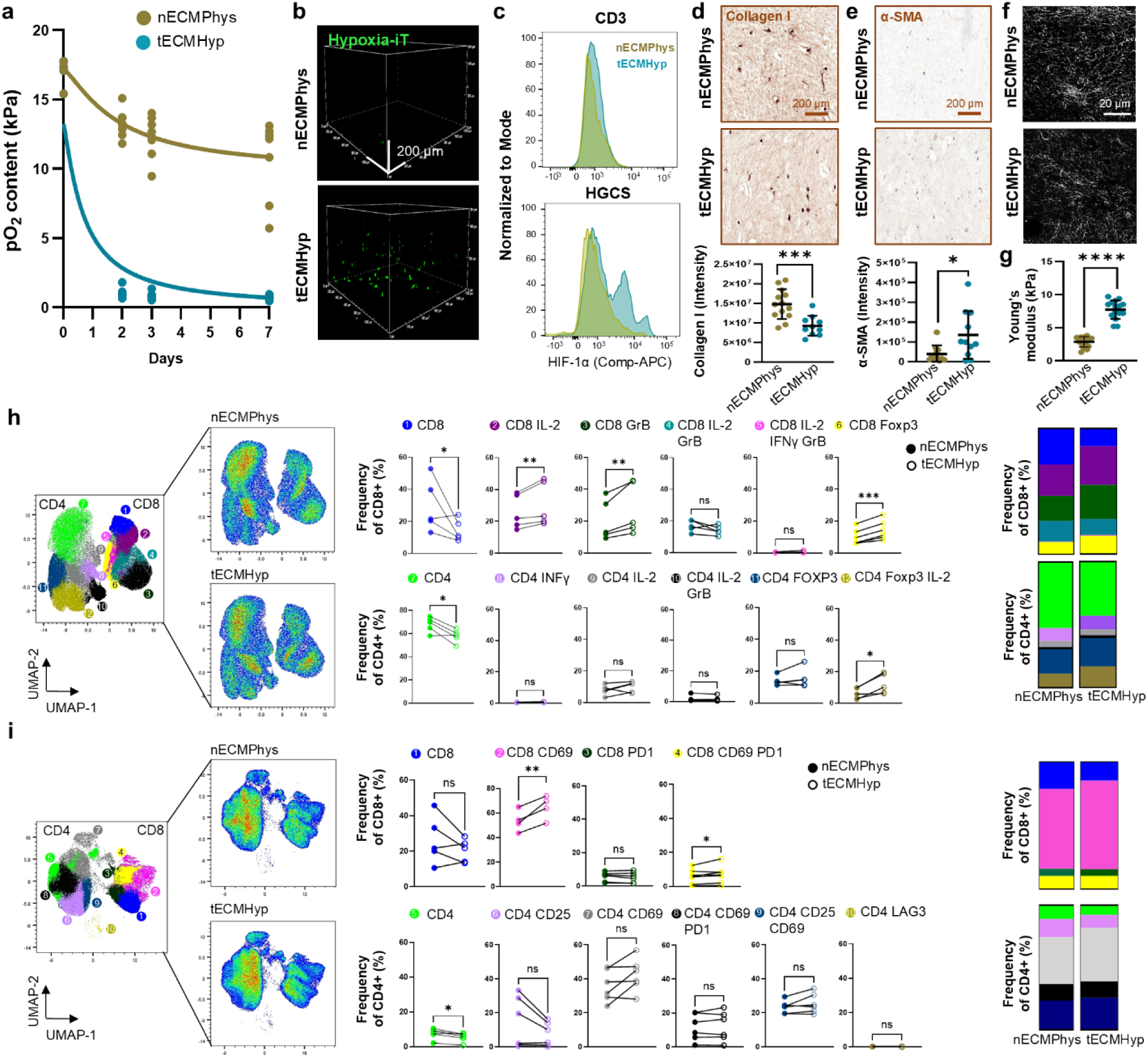
tECMHyp functionally recapitulates intratumoral hypoxia, ECM remodeling, and functionality of effector T cells. **a** Oxygen levels measured inside nECMPhys and tECMHyp models (0, 2, 3, and 7 days) by an optical sensor. **b** Confocal microscopy images in 3D view of hypoxic cells (Image- iT™ Green Hypoxia). z=500 μm, scale bar=200μm. **c** Histograms of HIF-1α expression in co-cultured primary HGSC tissue with autologous PBMCs. **d** IHC images for collagen I and quantification, scale=200μm. **e** IHC images for α-SMA and quantification, scale=200μm. **f** SHG images of collagen fibers in co-culture. Scale bar=20μm. **g** Stiffness quantification by AFM represented by Young’s Modulus (kPa). **h** UMAP plot, density plots, and quantification represented as frequency (percentage of parent CD4 or CD8) of intracellular markers. **i** UMAP, density plots, and quantification represented as frequency (percentage of parent CD4 or CD8) of activation and exhaustion surface markers. d, e, g Mean±SD, n=10-16, t-test; h-i, n=5, paired t-test. *p<0.05, **p<0.01, ***p<0.001,****p<0.0001, ns=not significant.

To explore the role of physoxia and hypoxia on ECM and the immune landscape, single-cell RNA sequencing (scRNA-seq) was performed in co-cultures of dissociated HGSC tumors with matching PBMCs in nECMPhys and tECMHyp using native PBMCs as a control. We identified seven single-cell clusters corresponding to different cell types in the TME (Fig. 1b,c and Extended Data Fig. 1a,b), including HGSC epithelial cells (HGSC), cancer-associated fibroblasts (CAFs), CD8+ and CD4+ lymphocytes (CD8 and CD4), natural killer cells (NK), B cells (B), and macrophages (Mφ). Immune cell type populations were identified in all experimental conditions (PBMCs, nECMPhys, and tECMHyp), while HGSC and CAFs cells were only present in nECMPhys and tECMHyp, as expected (Extended Data Fig. 1c). Marker genes were conserved in both tunable engineered models when compared to native PBMCs (Extended Data Fig. 1d), suggesting that both models largely capture the transcriptomic profile of immune populations present in PBMCs. Therefore, only nECMPhys and tECMHyp models were further compared to determine the influence of physiological and pathological oxygen levels in ECM dynamics and immune cell behavior. Classical hypoxia genes and hypoxic gene signatures correlated with bad prognosis in HGSC patients^31^ corroborated hypoxic features in tECMHyp when compared to nECMPhys (Fig. 1d and Extended Data Fig. 1e). Similarly, intratumoral hypoxia in tECMHyp influenced ECM remodeling at a transcriptomic level including enriched matrisome (ECM-related) genes, CAF-associated genes, and TGF-β pathway genes (Fig. 1e). While matrisome and CAF-associated genes were mostly exclusively expressed by CAFs, TGF-β signaling genes were expressed by all cell types (Extended Data Fig. 2a). To explore the effect of physiological oxygen levels on immune functionality, we evaluated activation, exhaustion and cytotoxicity genes and found them upregulated in tECMHyp when compared to nECMPhys, specifically expressed by cytotoxic effector cells including CD8+T and NK cells (Fig.1f and Extended Data Fig.2b). Moreover, when the transcriptional profile of nECMPhys and tECMHyp was compared, we found 11,695, 11,645, 11,212, 11,993, 10,368, 9,665, 11,395, and 8,927 differential expressed genes (DEG) for all populations, HGSC, CAFs, CD8, CD4, NK, B and Mφ, respectively (Extended Data Fig. 2c). Importantly, ECM remodeling (matrisome and TGF-signaling), as well as immune activation, exhaustion and cytotoxicity genes were significantly upregulated in tECMHyp compared to nECMPhys (Extended Data Fig. 2c and Supplementary Data 1). Furthermore, we investigated the probability and the strength of cancer-stroma-immune signaling interactions using CellChat. Stronger intercellular interactions were detected in tECMHyp compared to nECMPhys (Fig. 1g). A closer look at the specific interactions in each cell type with the rest of the cell populations confirmed that in tECMHyp there were stronger interactions of HGSC cells with CAFs, CD8+, and NK cells (Fig. 1h), and of CAFs with CD8+ and NK cells (Extended Data Fig. 3a) compared to nECMPhys. Similar trends were observed among other immune cells with NK and CD8+ T cells in the tECMHyp compared to nECMPhys and strong interactions of all cells with macrophages were observed independently of oxygen conditions (Extended Data Fig. 3a). To further explore the role of hypoxia and ECM remodeling on the cellular interactions in the TME, we studied the interactome of all cell types in the collagen pathway. The analysis of the strength and probability of the interactions of all cell populations in the collagen pathway indicated that CAFs are the main contributors with stronger interactions with CD8+ T cells, NK cells, B cells, and HGSC cells in tECMHyp when compared to nECMPhys, and also, they have a strong and high probability of interactions with Mφ independently of oxygen levels (Fig. 1i,j.). Under close examination, we observed that CAFs acted as the main senders, receivers, mediators, and influencers of the collagen pathway signals in both nECMPhys and tECMHyp confirming their key role in ECM remodeling and interactions with the rest of the cells through collagen signaling. Also, HGSC, CD8, and NK cells gained more roles as receivers of the signals in the collagen pathway in tECMHyp compared to nECMPhys (Extended Data Fig. 3b,c). Moreover, we identified important ligand-receptor pairs and their contribution in collagen pathway interactions such as the ligands (COL1A2, COL1A1, COL6A1) and receptors (CD44, SDC4, ITGA1+ITGB1, ITGA2+ITGB1, etc.) highlighting COL1A2-(ITGA1+ITGB1) and (ITGA2+ITGAB1) as the main contributors under intratumoral hypoxia (Extended Data Fig. 3d). Specifically, the probability of ligand-receptors interactions with the rest of the cells was higher in CAFs (Extended Data Fig. 3e) and in HGSC cells (Extended Data Fig. 3f) in the tECMHyp when compared to nECMPhys, even though there was a lower number of possible interactions correlated with collagen signaling pathway in the HGSC cells than in CAFs, indicating that CAFs are key players in ECM remodeling. Altogether, these data suggest that intratumoral hypoxia when compared to physoxia is a key driver of ECM remodeling through CAFs, sustains the functionality of cytotoxic effector cells via activation, exhaustion, and cytotoxicity, and influences cancer-stroma-immune interactions through collagen pathway, enabled by our physiologically relevant HGSC models.

### tECMHyp model functionally recapitulates intratumoral hypoxia, ECM remodeling, and functionality of effector T cells

To functionally characterize the transcriptional changes identified by scRNA-seq, we performed functional assays to validate intratumoral hypoxia, ECM remodeling, and immune functionality in the multi-cultures inside the 3D model recapitulating immune-inflamed or “hot” tumors when compared to a physoxic ovarian environment. The partial pressure of oxygen (pO_2_) inside the 3D scaffold was measured by an optical sensor and confirmed that the 3D models recreated physiologically relevant oxygen levels of HGSC-TME and the healthy ovary. When the 3D scaffold with the multi-culture of ovarian cancer cell lines, CAFs and PBMCs was incubated in 21% O_2_ incubator, the average oxygen levels inside the nECMPhys were approximately 12-13 kPa on days 2-3 and 10 kPa on day 7, recreating the physoxic O_2_ levels of healthy ovarian tissue. When incubated in a 1.5% O_2_ hypoxic chamber, the oxygen level in tECMHyp was significantly lower, reaching 0.8-1 kPa on days 2-3 and decreasing to 0.7 kPa on day 7 recreating the hypoxic HGSC-TME (Fig. 2a). Moreover, the hypoxic status of KURAMOCHI multi-cultured with CAFs and PBMCs inside of nECMPhys and tECMHyp was confirmed using a hypoxia-sensitive green dye. Green hypoxic cells were observed in multi-cultures (Fig. 2b) as well as monocultures (Extended Data Fig. 4a) in tECMHyp compared to nECMPhys. Furthermore, the expression of HIF-1α, a classical transcription factor overexpressed in hypoxia, was slightly higher in CD3+ T cells and robustly elevated in HGSC cells in tECMHyp compared to nECMPhys when primary HGSC tissue was co-cultured with autologous PBMCs and assessed by flow cytometry (Fig. 2c and Extended Data Fig. 4b). Similar results were observed in a multi-culture of KURAMOCHI with CAFs and PBMCs by IHC staining (Extended Data Fig. 4c). Therefore, this functional analysis revealed that nECMPhys and tECMHyp biomimetic models can recapitulate the physiologically relevant oxygen levels in the healthy ovary and the intratumoral hypoxia in HGSC-TME, respectively.

Second, to study oxygen-driven remodeling of the ECM, IHC staining revealed that a significantly higher expression of collagen I and a marker for CAFs activation, α-SMA, were observed in KURAMOCHI cells co-cultured with CAFs in tECMHyp compared to nECMPhys (Extended Data Fig. 4d,e), in the absence of PBMCs. When CAFs were grown in monoculture, there was also significantly higher expression of collagen I in tECMHyp compared to nECMPhys, even though it was two times lower than when they were co-cultured with KURAMOCHI, and no significant changes in α-SMA expression were observed. There was neither collagen I nor α-SMA detected in the KURAMOCHI monoculture (Extended Data Fig. 4d,e). When KURAMOCHI cells were multi-cultured with CAFs and PBMCs there was a significantly higher expression of α-SMA but lower expression of collagen I, although still higher compared to monocultures in the tECMHyp compared to nECMPhys (Fig. 2d,e). Considering the indirect relationship of collagen expression and matrix metalloproteinases (MMPs) activity, MMP-9 was investigated by ELISA and Western blot. MMP-9 was significantly higher in hypoxia compared to physoxia for multi-cultures containing PBMCs, but unaltered in monocultures and co-cultures of cancer and CAFs (Extended Data Fig. 4f,g and Source Data 1). These results confirm that CAFs are more activated and express higher levels of collagen in the hypoxic TME, while reflecting a dynamic response to microenvironmental cues, upregulated in co-culture with cancer cells but reduced upon the addition of immune cells, highlighting the context-dependent modulation of collagen production. Studies by second harmonic generation microscopy (SHG) in the co-culture of KURAMOCHI and CAFs confirmed that collagen fibers were detected only when the CAFs were present in the 3D scaffold independently of oxygen levels (Fig. 2f and Extended Data Fig. 4h). The orientation analysis revealed that the collagen fibers were highly aligned (Extended Data Fig. 4i). To further study how the increased collagen I secretion affected the biomechanical properties of the ECM, the stiffness of the nECMPhys and tECMHyp model when KURAMOCHI and CAFs were co-cultured inside the 3D scaffold was determined by atomic force microscopy (AFM) observing significantly higher stiffness of tECMHyp compared to nECMPhys (Fig. 2g). These results indicate that intratumoral hypoxia induces stiff aberrant ECM through collagen I secretion by activated CAFs and confirm that the 3D tECMHyp model mimics hypoxia-induced CAF-driven ECM remodeling when compared to physoxic normal ovarian ECM environment.

Third, we evaluated the impact of oxygen levels on immune-cancer interplay and immune effector functions. For that purpose, primary tumor cells from dissociated HGSC tumors were co-cultured with autologous PBMCs inside nECMPhys or tECMHyp made with matching plasma. Unbiased dimensionality reduction analysis using UMAP algorithm for an intracellular panel on T cells revealed that intratumoral hypoxia induced significantly higher activation (CD8+IL-2+) and cytotoxicity (CD8+GrB+) of CD8+ T cells and significantly higher frequency of T regulatory (T-reg) cells (CD4+Foxp3+IL2+ and CD8+Foxp3+) in tECMHyp compared to nECMPhys at the expense of reduced CD8+ and CD4+ T cells (Fig. 2h and Extended Data Fig.5a). Moreover, using a panel for activation and exhaustion surface markers on T cells, we identified that while the frequency of activated (CD8+CD69+) and exhausted (CD8+CD69+PD-1+) CD8+ T cells was significantly greater in tECMHyp compared to nECMPhys, CD4+ T cells only had a modest, not significant, trend to activation (CD4+CD69+) (Fig. 2i and Extended Data Fig. 5b). To functionally corroborate cytotoxic potential of the immune cells, apoptosis assays were performed in KURAMOCHI cells in monoculture or co-culture with PBMCs inside the 3D scaffold and the percentage of live, early apoptotic and late apoptotic/death cells was assessed by flow cytometry. While the KURAMOCHI apoptotic stage remained unaffected by oxygen conditions, the presence of PBMCs in co-culture with cancer in tECMHyp had a significant increase in apoptosis, both early and late apoptosis/death when compared to nECMPhys (Extended Data Fig. 5c). Furthermore, when PBMCs were co-cultured with KURAMOCHI at increasing PMBCs:HGSC ratios in 2D, we confirmed that the frequency of CD8+GrB+ increased in a concentration-dependent manner under hypoxia compared to normoxia, which translated into a higher cytotoxic effect in hypoxia by reducing significantly the percentage of live HGSC cells (Extended Data Fig. 5d). Taken together, these results indicate that intratumoral hypoxia induces higher effector functions in CD8+ cytotoxic T cells, by enhancing expression of activation and cytotoxicity markers, but also the activation of T-reg-like immunosuppressive cells in the tECMHyp model when compared to physiological nECMPhys.

### Hypoxia-induced CAF-driven ECM remodeling preserved in tECMHyp model causes impaired immune infiltration

Given that the majority of HGSC patients present immune-deserted or “cold” tumors characterized by immune exclusion^9^, we assessed the effects of hypoxia-induced CAF-driven ECM remodeling on immune infiltration. KURAMOCHI cells were monocultured or co-cultured with CAFs for three days to ensure physiologically relevant oxygen levels and allow ECM remodeling inside the nECMPhys or tECMHyp. On day 3, PBMCs were added on top and challenged to infiltrate into the 3D cultures and then evaluated by confocal microscopy and flow cytometry on day 7 (Fig. 3a). Co-cultures vs monocultures were established to delineate the individual effects of hypoxia vs hypoxia-induced CAFs. Confocal microscopy revealed a significantly decreased infiltration of CD45+ lymphocytes (green) and no significant changes to KURAMOCHI counts in tECMHyp compared to nECMPhys when HGSC was co-cultured with CAFs (Fig. 3b). While infiltrated CD45+ cells were also significantly reduced in tECMHyp compared to nECMPhys when HGSC were monocultured, HGSC counts were significantly increased (Extended Data Fig. 6a), suggesting competition for nutrients or altered metabolic support in hypoxic co-cultures. An immunophenotyping study identified eight populations including six immune cell populations (CD8+ T cytotoxic cells, CD4+ T helper cells, CD14+ Mφ, CD19+ B cells, CD56+ NK cells and CD11b+ MDSC), HGSC (B7H3+), and CAFs (FAP+) by dimensionality reduction analysis using UMAP algorithm (Fig. 3c and Extended Data Fig. 6b). Significantly impaired immune infiltration of CD8+ T cells, CD4+ T cells, B cells, Mφ, and MDSCs was observed in tECMHyp compared to nECMPhys while no significant changes were detected in NK cells, HGSC nor CAFs when KURAMOCHI was co-cultured with CAFs. When KURAMOCHI was monocultured, there was significantly impaired infiltration of Mφ in tECMHyp compared to nECMPhys, however no significant differences among the rest of the immune cell populations (Extended Data Fig. 6c). Similarly to confocal microscopy results, a significantly higher number of HGSC cells was detected in the tECMHyp compared to nECMPhys when cultured alone. These results reveal that the hypoxia-induced CAF-driven ECM remodeling preserved in the tECMHyp model causes impaired immune infiltration and further highlight the feed-forward loop effect of intratumoral hypoxia on CAFs influencing HGSC counts and immune evasion mechanisms.

**Fig. 3.**
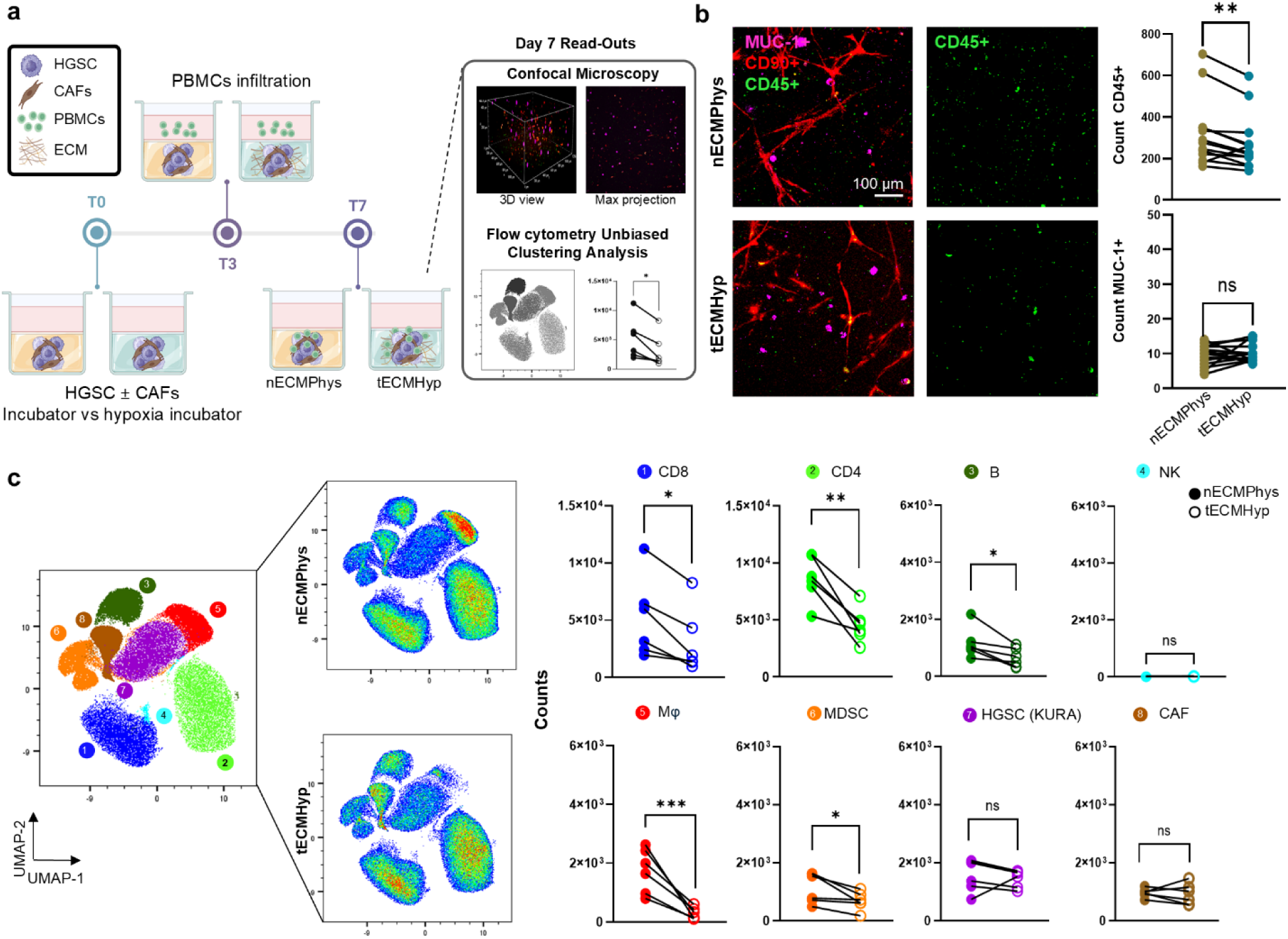
Hypoxia-induced CAF-driven ECM remodeling preserved in tECMHyp model causes impaired immune infiltration. **a** Schematic representation of methods used in immune infiltration studies. **b** Representative images of confocal microscopy z-stack view as maximum projection for CD45+ PBMCs (green) infiltrated into the scaffold with HGSC (MUC-1+, magenta) and CAFs (CD90+, red) and quantification of number of infiltrated CD45+ and resident MUC-1+ HGSC cells. Scale bar=100μm. n=14. **c** UMAP plot, density plots, and quantification of number (counts) of infiltrated PBMCs (CD3+, CD4+, and CD8+ T cells, Mφ (CD14+), NK (CD56+), B (CD19+) and MDSC (CD11b+)), and HGSC and CAFs cultured inside the 3D scaffolds using dimensionality reduction analysis by flow cytometry. n=6. Paired t-test, *p<0.05, **p<0.01,***p<0.001, ns=not significant.

### Preincubation with galunisertib reduces hypoxia-induced CAF-driven TGF-β signaling and collagen I expression recapitulated in tECMhyp model

In CAFs, TGF-β signaling plays a crucial role in driving collagen I secretion and ECM remodeling^32, 33^. Intratumoral hypoxia upregulates TGF-β and its receptors, leading to increased SMAD activation, promoting ECM remodeling through activation of CAFs, stimulating the expression of collagen (COL1) and lysyl oxidase (LOX) genes, and repression of MMP genes, aiding immune escape^32, 34^. To prevent the hypoxia-induced dysregulated ECM remodeling involved in impaired immune infiltration, we pharmacologically targeted TGF-β signaling using preincubation with galunisertib (Gal), a small-molecule inhibitor of the TGF-β receptor I kinase (Fig. 4a). Higher expression of TGF-β in tECMHyp compared to nECMPhys was validated by western blot in co-cultures (Fig. 4b and Source Data 2a), which aligned with the transcriptomic data from Fig. 1e. When monocultures were interrogated, it was found that CAFs were primarily responsible of the TGF-β upregulation (Extended Data Fig. 7a and Source Data 2a). Pre-incubation with Gal induced downregulation of downstream canonical signaling (TGF-β, pSMAD2/3, SMAD4) and non-canonical pathways (pMAPK and pAKT) when compared to untreated control in both nECMPhys and tECMHyp in co-cultures (Fig. 4c and Source Data 2b) and more strongly in the CAFs than the HGSCs cells in monocultures (Extended Data Fig. 7b and Source Data 2b). Intracellular staining using imaging flow cytometry quantitatively validated that pre-incubation with Gal significantly reduced TGF-β expression in both nECMPhys and tECMHyp in co-cultures and proved CAFs as the main contributors to intracellular TGF-β expression (Fig. 4d and Extended Data Fig. 7c). Monocultures further supported CAFs robust implication in intracellular TGF-β expression with significant downregulation by Gal treatment (Extended Data Fig. 7d). Of note, a significantly increased intracellular TGF-β expression was identified by Gal treatment in nECMPhys in opposition to a reduced expression in tECMHyp, but still HGSC expression was significantly lower than in CAFs.

**Fig. 4.**
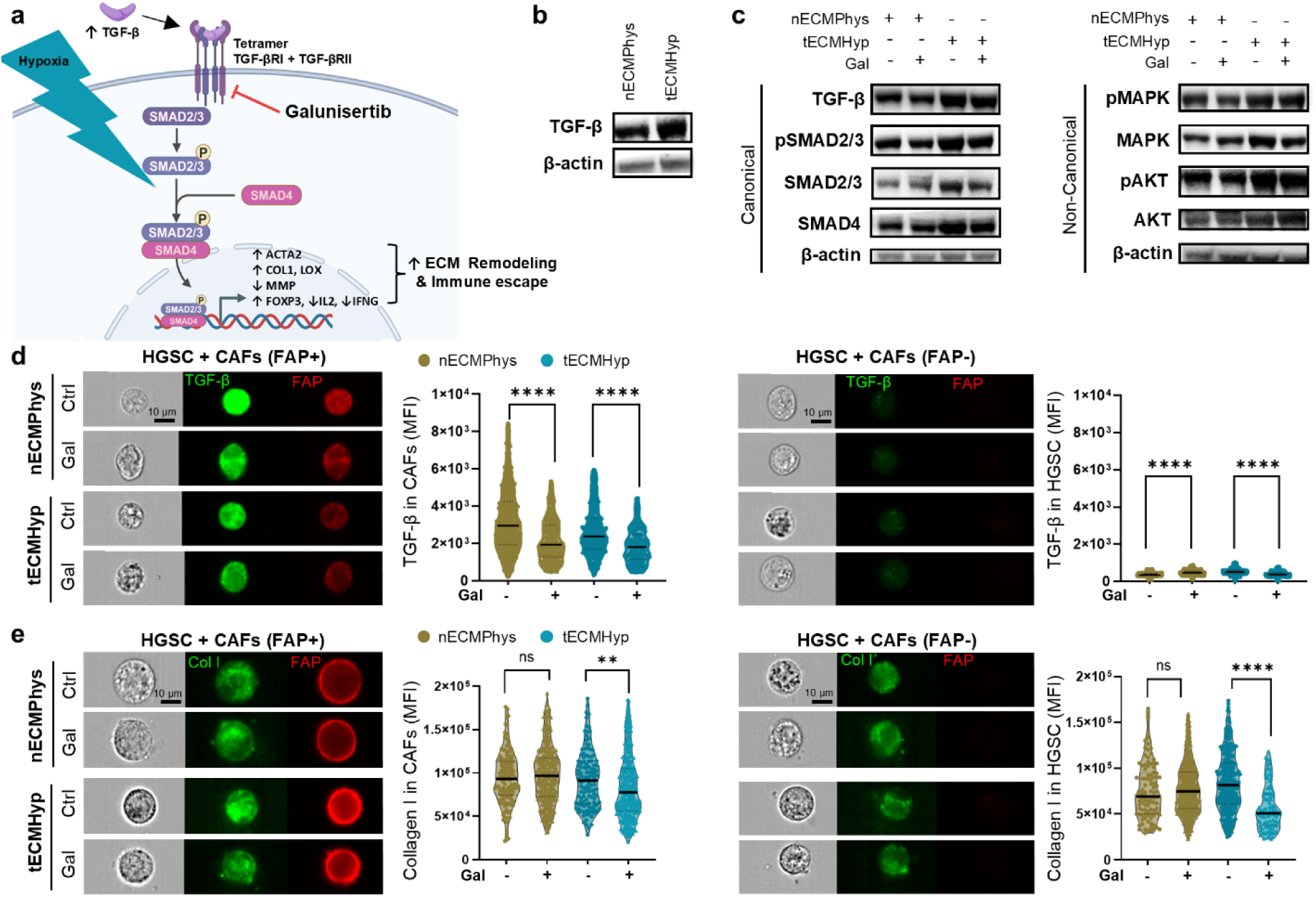
Preincubation with galunisertib reduces hypoxia-induced CAF-driven TGF-β signaling and collagen I expression recapitulated in tECMHyp model. **a** Schematic of TGF-β signaling pathway involved in ECM remodeling in the hypoxic TME and the mechanism of action of TGF-βRI inhibitor, galunisertib. **b** Representative images of TGF-β expression detected by western blot in co-culture. **c** Expression of signaling molecules involved in canonical (left) and non-canonical (right) TGF-β signaling pathway in the presence or absence of galunisertib (Gal) detected by western blot in co-culture. **d** Representative images of intracellular TGF-β (green) and **e** collagen I (green) in imaging flow cytometry in co-culture of CAFs (FAP+, red) and HGSC (FAP-), and quantification of green mean fluorescence intensity (MFI). Scale=10 μm. Median. Mann-Whitney test, **p<0.01, ****p<0.0001, ns=not significant.

In light of the relationship between TGF-β and collagen type I, we ascertained whether galunisertib treatment affected collagen expression and its deposition. Intracellular collagen staining aligned with the results of intracellular TGF-β expression where Gal significantly decreased intracellular collagen in tECMHyp when compared to untreated control for both CAFs and HGSC in monoculture and co-culture (Fig. 4e and Extended Data Fig. 7e). Furthermore, cellular and secreted collagen type I were evaluated by IHC staining and confirmed that pre-incubation with galunisertib significantly reduced collagen expression in tECMHyp in CAFs monocultures and their co-cultures with KURAMOCHI, however, increased in nECMPhys co-cultures (Extended Data Fig.7f). Collectively, our results demonstrate that preincubation with galunisertib reduces robustly the hypoxia-induced CAF-driven TGF-β signaling and collagen I expression recapitulated in tECMHyp models, opening ways to normalize the dysregulated ECM remodeling in HGSC tumors.

### Preincubation with galunisertib rescued the hypoxia-induced CAF-driven impaired immune infiltration

To further explore if galunisertib could reverse the hypoxia-induced CAF-driven impaired immune infiltration, nECMPhys and tECMHyp models were preincubated for three days in the absence or presence of galunisertib and infiltration studies were performed as previously discussed (Extended Data Fig.8a). Preincubation with galunisertib significantly rescued the impaired infiltration of CD8+ T cells, CD4+ T cells, and B cells in tECMHyp compared to not treated control, and no significant changes were observed in the rest of the immune cell types or in the physoxic conditions when autologous PBMCs were challenged to infiltrate into the 3D matrix with primary HGSC cells from dissociated tumors co-cultured with CAFs (Fig. 5a and Extended Data Fig. 8b). Of note, a broader spread of UMAP clusters was identified when using autologous PBMCs from HGSC patients revealing underlying biological variability or inter-patient heterogeneity that could be related to variable treatment response or divergence in cellular landscape (Table 1). Analysis of the secreted tumor milieu revealed a higher expression of pro-tumoral cytokines such as IL-1β and growth factors such are GM-CSF and VEGF-A in tECMHyp compared to nECMPhys and no significant changes in anti-tumoral cytokines based on the oxygen conditions except for IL-18 (Fig.5b). Preincubation with galunisertib decreased the expression of pro-tumoral growth factors (G-CSF, GM-CSF, and VEGF-A) in nECMPhys and tECMHyp, but also enhanced pro-tumoral cytokines such are IL-10 and MIP-1β and IL-18.

**Fig. 5.**
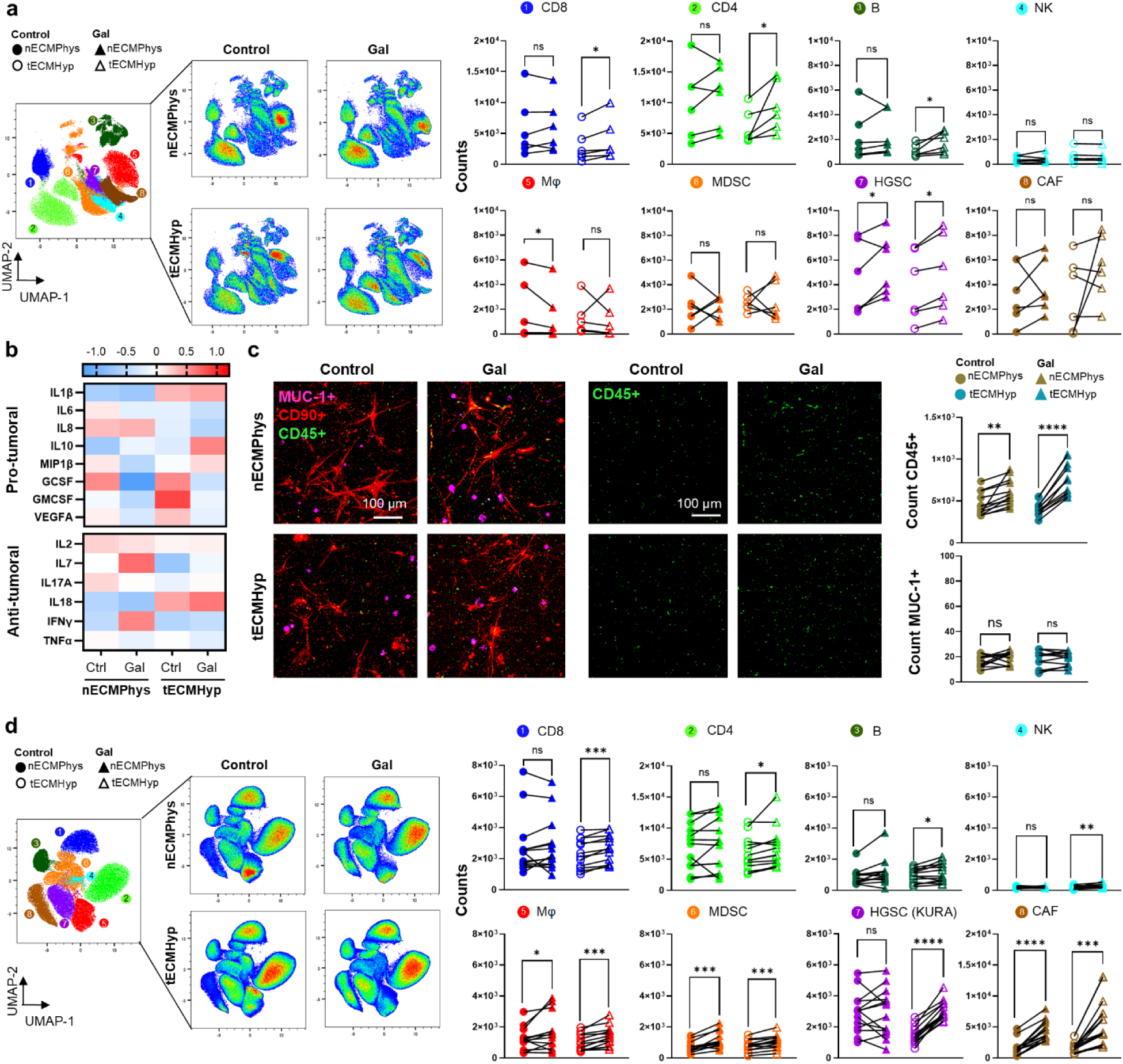
Preincubation with galunisertib rescued the hypoxia-induced CAF-driven impaired immune infiltration. **a** UMAP plot, density plots, and quantification of the number (counts) of infiltrated PBMCs cell types (CD3+, CD4+, CD8+ T cells, Mφ (CD14+), NK (CD56+), B (CD19+) and MDSC (CD11b+)), and HGSC cells from dissociated tumors in co-culture with CAFs inside the 3D models in the presence or absence of galunisertib (Gal) in dimensionality reduction analysis by flow cytometry. n=6. **b** Heatmaps showing the expression of secreted pro-tumoral and anti-tumoral cytokines. Median of z-score for each analyte. n=6. **c** Confocal microscopy images of z-stack view as a maximum projection of infiltration of PBMCs (CD45+, green) into the scaffold with KURAMOCHI (MUC-1+, magenta) and CAFs (CD90+, red) in the presence or absence of galunisertib and quantification of the number of CD45+ and HGSC. Scale=100μm. n=12. **d** UMAP, density plots, and quantification of number (counts) of infiltrated PBMCs phenotypes and KURAMOCHI and CAFs inside the scaffold in the presence or absence of galunisertib in dimensionality reduction analysis by flow cytometry. n=13. Paired t-test, *p<0.05, **p<0.01, ***p<0.001, ****p<0.0001, ns=not significant.

**Table 1.**
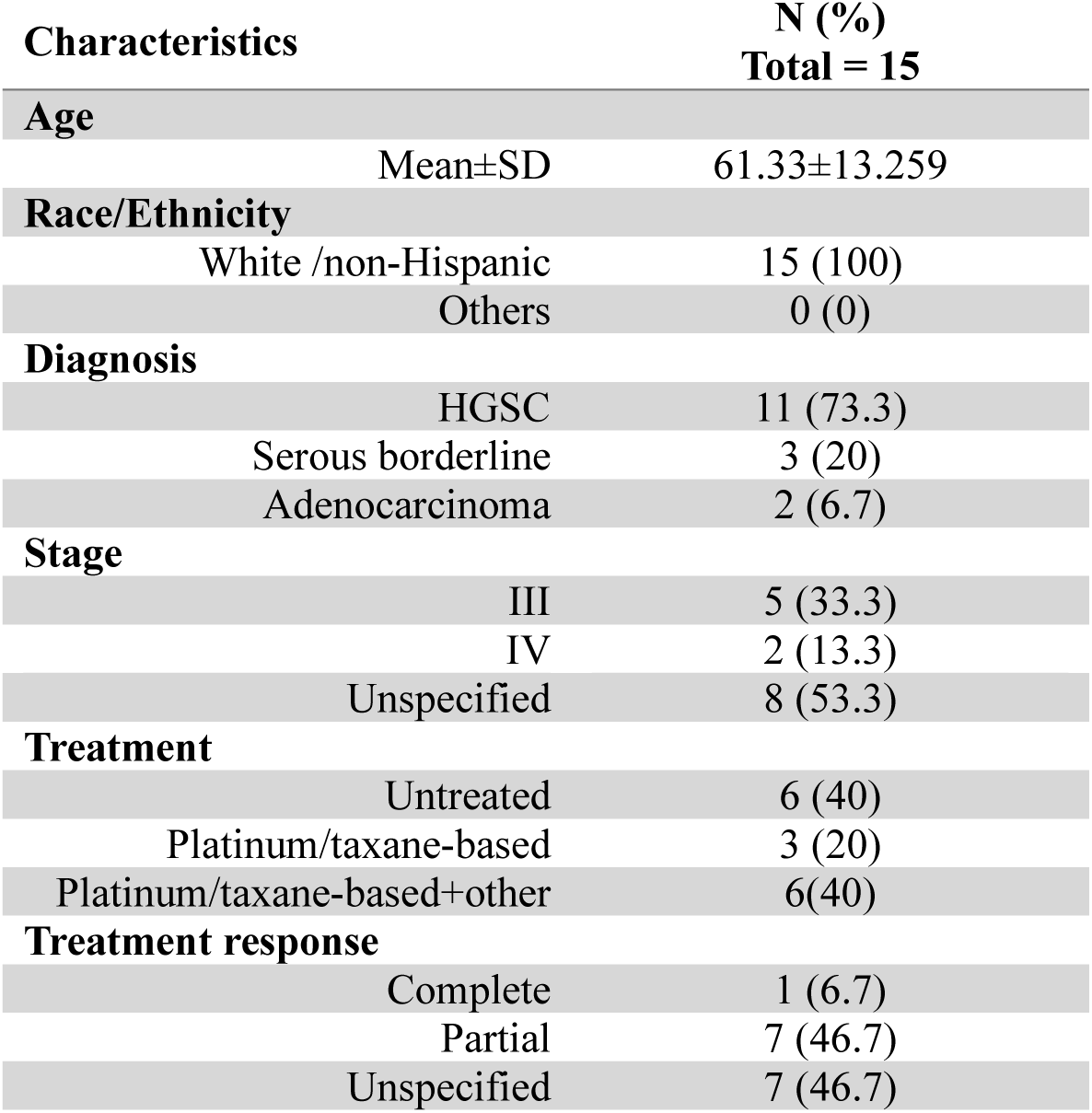
Demographics of patients used in the study.

| Characteristics |  | N (%)<br>Total = 15 |
| --- | --- | --- |
| **Age** |  |  |
|  | Mean±SD | 61.33±13.259 |
| **Race/Ethnicity** |  |  |
|  | White /non-Hispanic | 15 (100) |
|  | Others | 0 (0) |
| **Diagnosis** |  |  |
|  | HGSC | 11 (73.3) |
|  | Serous borderline | 3 (20) |
|  | Adenocarcinoma | 2 (6.7) |
| **Stage** |  |  |
|  | III | 5 (33.3) |
|  | IV | 2 (13.3) |
|  | Unspecified | 8 (53.3) |
| **Treatment** |  |  |
|  | Untreated | 6 (40) |
|  | Platinum/taxane-based | 3 (20) |
|  | Platinum/taxane-based+other | 6(40) |
| **Treatment response** |  |  |
|  | Complete | 1 (6.7) |
|  | Partial | 7 (46.7) |
|  | Unspecified | 7 (46.7) |

Complementary experiments were performed in the KURAMOCHI cell line co-cultured with CAFs and infiltrating PBMCs to validate the observed effects of galunisertib in immune infiltration while minimizing confounding effects of inter-patient heterogeneity by confocal imaging and flow cytometry. Confocal microscopy results showed that preincubation with Gal significantly increased immune infiltration into the 3D matrix in tECMHyp and nECMPhys compared to untreated controls and no significant changes were observed in the number of KURAMOCHI cells co-cultured with CAFs (Fig.5c). A similar increase in CD45+ T cells infiltration was detected in KURAMOCHI monocultures, accompanied by higher cancer counts in the tECMHyp preincubated with galunisertib compared to control (Extended Data Fig. 9a). Immunophenotyping results corroborated that galunisertib significantly rescued the impaired immune infiltration of CD8+ T cells, CD4+ T cells, B cells, NK cell, Mφ, and MDSC compared to untreated controls, along with a significantly increased number of HGSC and CAFs in the tECMHyp treated compared to control (Fig.5d and Extended Data Fig. 9b). When KURAMOCHI was monocultured, there was a significantly increased immune infiltration of CD8+ T cells, CD4+ T cells, B cells, NK cells, and Mφ in the tECMHyp pre-treated with Gal compared to control, but no significant changes in MDSCs and KURAMOCHI cells (Extended Data Fig. 9c). To further determine whether the rescued immune infiltration by galunisertib in hypoxia aligns with an increased cytotoxic effect, confocal microscopy with live/dead cell staining was performed and verified a significantly higher cell death in the presence of Gal compared to control (Extended Data Fig. 9d). In general, galunisertib reversed the hypoxia-induced impaired infiltration of lymphoid lineage cells (CD8+ T cells, CD4+ T cells, and B cells) with context-dependent effects in cancer, CAFs and myeloid lineage cells.

### Validation of galunisertib reversal of impaired immune infiltration in ovarian native dECM exposed to physiologically relevant oxygen levels

Acknowledging that the tunable patient-derived 3D models based on fibrin allow for hypoxia-induced CAF-driven ECM remodeling, but lack native biochemical and biomechanical ECM properties, we employed decellularized ovary matrices re-cellularized with KURAMOCHI and CAFs while exposed to physiologically relevant oxygen levels and then challenged PBMCs to infiltration, to further validate and support whether our previous findings are translationally relevant (Fig. 6a). Decellularization was confirmed by gross morphological assessment where decellularized tissue became translucent and whitish, consistent with removal of cellular material (Fig. 6b). Histological assessment included confirmation of loss of nuclei by H&E staining and preservation of ECM by Masson’s trichrome, collagen I and fibronectin staining in the decellularized matrix compared to native tissue (Fig. 6c). After corroboration of cell removal and ECM integrity, we validated that physiological oxygen levels and ECM remodeling were mimicked in these models at day 3. The partial pressure of oxygen (pO_2_) inside the re-cellularized sectioned tissue revealed approximately 16.3 kPa and 1.5 kPa levels when grown in 21% and 1.5% O_2,_ respectively, and significantly higher collagen I expression in hypoxia when compared to physoxia (Fig. 6d,e). Therefore, we refer to them as, normal decellularized extracellular matrix in physoxia (ndECMPhys) and tumorous decellularized ECM in hypoxia (tdECMHyp). We further verified by confocal microscopy that when immune cells were added onto these models, a significantly impaired immune infiltration in tdECMHyp compared to ndECMPhys was found (Fig. 6f), consistent with the results of the tunable patient-derived 3D models in Fig. 3. Then, preincubation with galunisertib significantly reversed the hypoxia-induced CAF-driven impaired infiltration of PBMCs in alignment with our previous results (Fig. 6g).

**Fig. 6.**
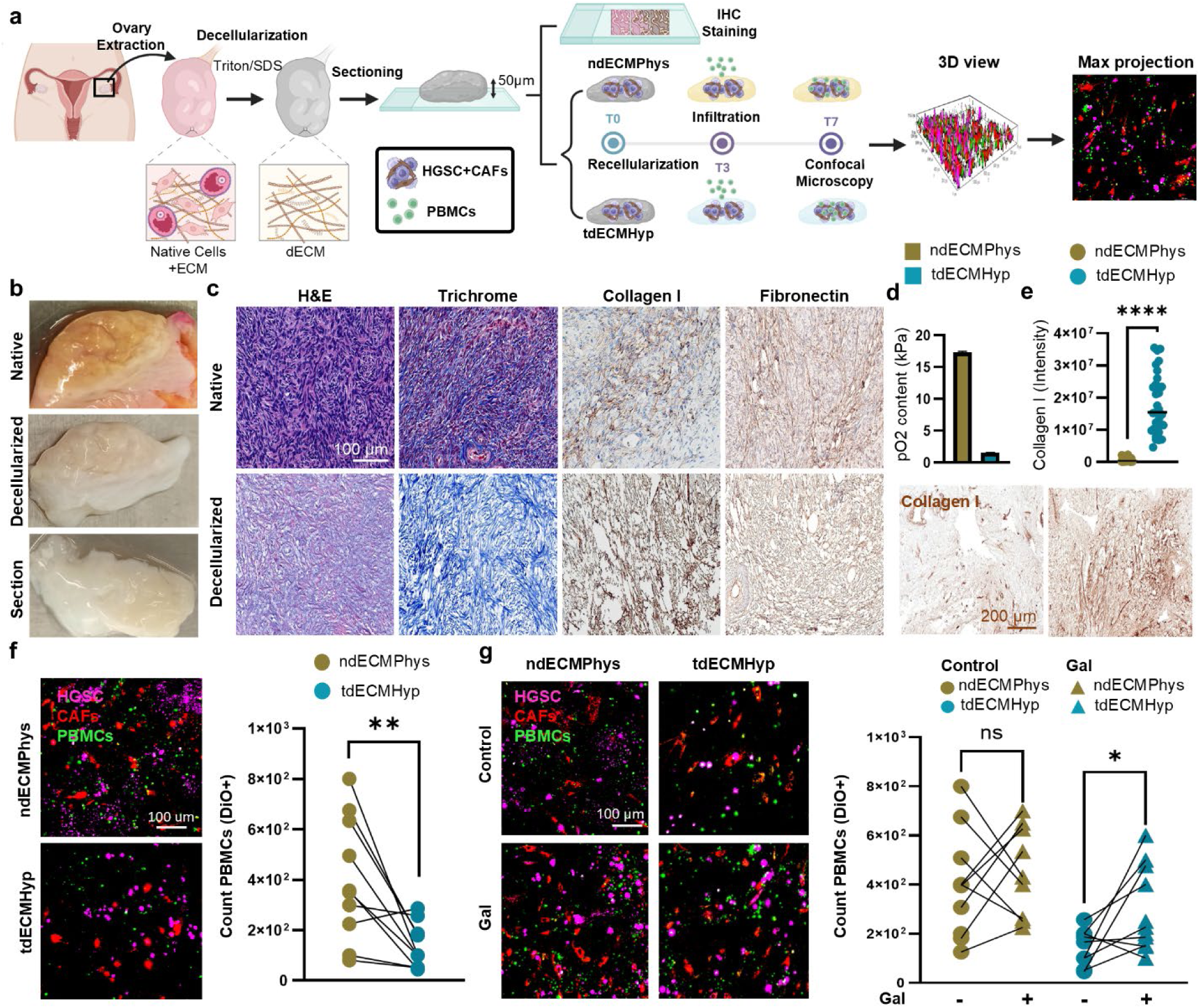
Galunisertib reverses hypoxia-induced CAF-driven impaired immune infiltration in ovarian native dECM. **a** Schematic representation of decellularization of human ovaries and subsequent recellularization in ndECMPhys (normal decellularized ECM Physoxia) and tdECMHyp (tumorous decellularized ECM Hypoxia) and immune infiltration studies. **b** Representative images of the human ovary before (top) and after decellularization (middle) and transversal section of dECM (bottom). **c** Representative images of H&E, Masson’s trichrome, collagen I, and fibronectin staining of ovarian native tissue and dECM. Scale=100μm. **d** Oxygen levels measured inside the re-cellularized ndECMPhys and tdECMHyp model by an optical sensor at day 3. n=4 **e** Quantification and IHC images for collagen I in re-cellularized dECM at day 3, scale=200μm. **f** Confocal microscopy images and the quantification of the number of infiltrated PBMCs (green) into the dECM recellularized with KURAMOCHI (magenta) and CAFs (red) in the absence and **g** in the presence of galunisertib. Scale=100μm. e, n=30, t-test; f-g, n=11,paired t-test, *p<0.05, **, p<0.01, ****p<0.0001 ns=not significant.

In summary, our findings show that the hypoxia-induced CAF-driven impaired immune infiltration can be recapitulated in the patient-derived tumor-immune tunable models, which were also suitable for dissecting targeted opportunities via TGF-β signaling and collagen by galunisertib, turning immunologically cold HGSC tumors into hot ones.

## Discussion

We bioengineered patient-derived tumor-immune tunable models that provide translationally relevant platforms to study tumor-stroma-immune dynamics in physio- and pathophysiologically relevant environments and overcome the limitations of current preclinical immune models which lack physiologically relevant tumor characteristics in terms of oxygen levels and ECM dynamics. Our approach relies on models that have been bioengineered to recapitulate physiologically relevant oxygen levels of hypoxic tumors and physoxic ovaries and hypoxia-induced ECM remodeling. These models integrate cancer cells co-cultured with cancer-associated fibroblasts (CAFs) exposed to immune cells as multi-culture or challenged to infiltrate into a 3D model bioengineered with autologous patient’s plasma or onto decellularized human ovaries. This study demonstrates that hypoxia-induced CAF-driven ECM remodeling impairs immune infiltration, which is in line with the literature^35–38^, while using physoxic ovarian environments as a better control for modeling the gradient between normal tissue and hypoxic tumors.

Tumors initiate in physoxic regions and progress into hypoxia while remaining surrounded by healthy tissue^39^, which has been frequently overlooked *in vitro* in favor of atmospheric oxygen levels typically used in standard cell culture^40–42^. Tumors are also characterized by an aberrant stiffer ECM, with a distinct composition from normal tissue^43^. When cells are cultured on or within ECM with composition or abundance that do not reflect their native or disease-specific environment, they can exhibit artificial or misleading behaviors^16, 17, 19^. Incorporating these physiologically relevant characteristics into *in vitro* models is essential to faithfully uncover context-specific mechanisms driving immune evasion^44, 45^. As an example of the relevance of physoxic oxygen levels in the ovaries, it has been demonstrated that lower oxygen tensions in *in vitro* culture yield higher follicle viability and quality than 20% O_2_ tension^46^. Matrix-based platforms such as collagen or basement membrane extracts have enabled the evaluation of tumor invasion and screening of therapies^37, 47, 48^. However, they neglect either the incorporation of stromal or immune compartments, or oxygen–induced shifts in matrix composition and immune cell distribution or inaccurately compare hypoxia to normoxia. Decellularized omental metastasis importantly demonstrated that the ECM educates an immunoregulatory tumor macrophage phenotype^19^, but this study was performed under a normoxic environment. To the best of our knowledge, our human-relevant tumor-immune tunable models are the first HGSC models recapitulating pathophysiological oxygen levels of the hypoxic HGSC-TME in comparison to the physoxic ovary environment, while incorporating CAFs and patient-derived materials such as tumor biopsy, PBMCs, and plasma. Critically, mixing patient-derived plasma with collagen to grow tumors enhances the physiological relevance of our *in vitro* models by incorporating systemic, patient-specific factors such as cytokines, metabolites, and growth factors^29, 30^. This approach supports precision-based applications by better capturing inter-individual variability in tumor behavior and therapeutic response^49^. Alternatively, to account for native ECM composition and architecture while reducing intra-assay variability due to the plasma matrix, decellularized ovaries were used as a validation platform, highlighting the important contribution of oxygen levels and CAF-driven ECM remodeling to immune evasion. Overall, our models are versatile tools that could be used to explore other cancer types as they are easily modifiable to recreate specific biological, biophysical, and biomechanical properties of other organs^29, 30^.

Mechanistically, we identified that the tECMHyp model captures classical hypoxic features at transcriptomic levels and via functional read-outs. We functionally characterize that intratumoral hypoxia induced CAF activation leading to aberrant ECM remodeling through overexpression of collagen and other core matrisome genes driven by TGF-β signaling, and densely packed and highly aligned collagen fibers with stiffer matrix mimicking key features of HGSC tumors^43, 44^. Other studies also supported that hypoxia induces collagen deposition in human and murine omental metastasis^50^. Remarkably, the presence of PBMCs in the multi-culture with cancer cells and CAFs, decreased the hypoxia-induced CAF-driven collagen I expression which correlated with an increased expression of MMP-9 similar to studies in other cancers^51, 52^, highlighting the importance of tumor-stroma-immune interactions^51^ in the hypoxic environment. Of note, we used fully characterized reprogrammed HGSC-CAFs generated in our lab^53^ that resemble primary CAFs while exhibiting high CAF and ECM/matrisome-scoring and were linked to the mesenchymal subtype with the worst prognosis in ovarian cancer^54^. Future studies should confirm the results using primary CAFs, bearing in mind they constitute a heterogeneous population that can lead to pleiotropic responses and present limited cell passage number and growth rate^55^. Based on functional measurements, we have shown that intratumoral hypoxia acts as a friend and foe in modulating immune responses, increasing the effector functions of CD8+ T cells with enhanced activation, cytotoxicity, and killing, while simultaneously promoting immunosuppressive cells. Similar findings were observed in other studies in HGSC patients where intratumoral hypoxia induced immunosuppressive cell types^56, 57^, but also a higher presence of activated CD8+ T cells in the TME^21, 58, 59^. TILs, particularly CD8+ T cells, existed along a functional spectrum simultaneously displaying features of activation, cytotoxicity, and exhaustion, reflecting a dynamic balance between ongoing tumor engagement and progressive dysfunction, which has been featured as a state predictive of immunotherapy response^60, 61^. In recent years, CAFs are emerging as important regulators of the immune cell continuum inducing immunosuppression and ovarian cancer progression^62, 63^. We showed that intratumoral hypoxia influenced cancer-stroma-immune interactions through the collagen pathway where CAFs were the main contributors in line with other studies^50, 63^. Strong communication probabilities were detected between the collagen I ligand and integrins, CD44, and syndecan receptors, which have been previously correlated with an immunosuppressive TME and chemoresistance in different cancers including ovarian cancer^63–66^. Infiltration studies demonstrated that intratumoral hypoxia alone is not responsible for immune evasion but rather it triggers CAF-driven ECM remodeling, which altogether impairs immune infiltration. This highlights the complexity of the TME and the need for sophisticated models capturing the feed-forward loop between oxygen content and CAF activation and its influence on immune dynamics.

Hypoxia induced the activation of CAFs leading to increased collagen and TGF-β expression, confirming CAFs as main orchestrators of TGF-β signaling and its correlation with hypoxia through HIF-1α as previously described^32^. Several strategies have been developed to target TGF-β signaling to overcome ECM remodeling and immune resistance^67^. Here, we used Galunisertib (Gal) as a model drug, which is a small molecule inhibitor of TGFβRI phosphokinase activity^68, 69^ and has been used in several preclinical models and clinical trials with promising results in different cancers such as hepatocellular carcinoma, myelodysplastic syndrome, rectal, and NSCLC^67–69^; however, limited efficacy in recurrent glioblastoma^70^ and metastatic pancreatic cancer^71^. We selected Gal both due to its reduced anti-tumor activity and simultaneous effects on CAFs and immune cells by reducing CAF activation and antifibrotic effects, and alleviation of immune suppression with higher infiltration and activation^69, 72, 73^. We acknowledge that Gal can present several side effects such as cardiotoxicity and skin toxicity, but they could be overcome by careful dosage, pulsatile therapy or targeted therapy^74^. We mechanistically confirmed the anti-fibrotic potential of Gal decreasing the expression of TGF-β signaling molecules and collagen I in hypoxia-induced CAFs similar to others^69, 75, 76^. Gal preincubation might also be weakening tumor supporting signals particularly relevant to myeloid recruitment. The treatment with Gal in physoxia induced higher expression of TGF-β in HGSC and collagen I in their co-cultures with CAFs. We hypothesize that Gal produces higher inhibition of TGF-β signaling correlated to collagen I synthesis in hypoxia where there is synergistic overexpression of HIF-1α and TGF-β, meanwhile, in physoxia it triggers compensatory mechanisms.

Importantly, our data indicates that pre-treatment with Gal rescued the hypoxia-induced CAF-driven impaired immune infiltration in both our plasma 3D models and decellularized ovaries, mainly the lymphoid lineage with context-dependent effects in myeloid lineage, parallel to other studies^77, 78^.

We acknowledge that our studies might have certain limitations. We investigated the role of hypoxia in inducing ECM remodeling, but we did not address how ECM remodeling affects hypoxia, even though their relationship is reciprocal^79^. The use of non-HLA or donor-matched PBMCs and CAFs may present a potential concern with alloreactivity, but due to the short duration window of the *in vitro* experiments (3 – 7 days) we expect minor signs of graft versus host disease, considering those have been identified within 4 weeks of PBMC implantation in humanized mice^80, 81^. Therefore, we mitigated this concern by levering autologous PBMCs and tumor cells from HGSC patients in part of our studies with the 3D models made with matching plasma to avoid the graft-vs-tumor rejection effects^82^. The differences in the responses between PBMCs from healthy donors and HGSC patients can be correlated with acquired immune resistance in primary cells and an altered cell behavior due to previous chemotherapeutic and immunotherapeutic clinical exposure in the HGSC patients^83^. Moreover, while the HGSC patient samples share some global transcription features, their internal diversity leads to more diffuse projections in low-dimensional space. We note that all participants were White/Caucasian non-Hispanic women, due to higher incidence and sample availability; however, samples from diverse patients should be studied to consider the race/ethnicity factor and prevent health disparities^84^. Another limitation is the lack of spatio-temporal organization of the cells inside of the 3D model through compartmentalization, which could be implemented using a tumor-on-a chip as previously described^85^. We also acknowledge that the physoxic O_2_ level in the dECM was higher than in normal tissues due to the low thickness of the tissue for imaging purposes, therefore increasing the thickness of the dECM or using an incubator with controlled O_2_ levels could better recapitulate physoxic levels in this model. In the future, more stromal cell types, such are endothelial cells or adipocytes, should be incorporated into the model to better mimic HGSC-TME. Further studies should also investigate the polarization of tumor-associated macrophages (TAMs) to decipher specifically how intratumoral hypoxia affects the innate immune response. Furthermore, different ECM-targeting drugs, or combination therapy of Gal, immunotherapy, and hypoxia-targeting agents should be explored but it was outside of the scope of the current study^67, 86^.

In summary, we have shown that recapitulation of the hypoxic tumor levels in comparison to the physoxic ovarian oxygen levels as a more physiologically relevant control are essential to gain insights into tumor-stroma-immune interactions and model immunologically cold HGSC tumors. Our findings underscore the pivotal role of hypoxia in driving CAF-driven ECM remodeling, which contributes to immune exclusion sustaining the immunologically cold nature of HGSC tumors. Importantly, our translationally relevant models offer a valuable tool for identifying novel therapeutic targets aimed at disrupting stromal barriers and promoting immune infiltration, offering a path toward transforming immunologically cold tumors into hot and responsive ones.

## Methods

### Cell lines and cell culture

KURAMOCHI (RRID:CVCL1345), OVSAHO (RRID:CVCL_3114), OVCAR-3 (ATCC HTB-161), OVCAR-8 (RRID:CVCL_1629), CAOV3 (ATCC HTB-75), OV81.2 and HEYA8 (RRID:CVCL_8878) cell lines were used as models for high-grade serous ovarian carcinoma and cultured in DMEM (MT10013CV, Corning)/Ham’s F-12 (MT10080CV, Corning) media (1:1) supplemented by 10% (V/V) of fetal bovine serum (FBS, Gibco), 100 U/ml penicillin, and 100 mg/ml streptomycin (Corning CellGro). OVCAR-3, OVCAR-8, CAOV-3, HEYA-8, and OV81.2 cell lines were a kind gift from Dr. Analisa Difeo and OV81.2 was generated from HGSC patient’s ascites re-engrafted into immunodeficient mice^87^. Ovarian cancer-associated fibroblasts (CAFs) were previously developed and characterized in our lab by reprogramming primary human uterine fibroblasts (ATCC PCS-460-010) with conditioned media from KURAMOCHI^53^.

### Human tissue and primary cells

Tumor biopsy and peripheral blood samples from ovarian cancer patients undergoing a cytoreductive surgery were provided by the Sanford USD Medical Center, Sioux Falls, SD, USA. Informed consent was signed by all subjects with approval from the Sanford Health Institutional Review Board and in accordance with the Declaration of Helsinki (STUDY00001315). Peripheral blood from de-identified healthy female volunteers was provided by the Sanford Blood Biobank. Ovarian tumor tissue was stored in MACS® tissue storage solution (130100008, Miltenyi Biotec) and used fresh or frozen in freezing solution composed by FBS and 10% (V/V) dimethyl sulfoxide (DMSO, D8418, Sigma-Aldrich). Patient demographic and therapeutic treatment characteristics are included in Table 1. Tumor tissues were dissociated using MACS® Tissue Dissociation kit (130095929, Miltenyi Biotec) following the manufacturer’s instructions. Primary cells were cultured in DMEM/Ham’s F-12 (1:1) supplemented with 10% of FBS and 100 U/ml penicillin and 100 μg/ml of streptomycin. Peripheral blood mononuclear cells (PBMCs) were isolated from peripheral blood using Leucosep round bottom centrifugation tubes (07000500, Greiner Bio-One, Fisher Scientific) filled with Ficoll®-Paque Plus density gradient media (GE17444003, Sigma-Aldrich). PBMCs were cultured in CTS™ OpTmizer™ T Cell Expansion SFM (A1048501, Gibco) supplemented with OpTmizer™ T-Cell Expansion Supplement and 2 mM L-glutamine (Gibco), and if needed, stimulated in RPMI media with eBioscience™ Cell Stimulation Cocktail (plus protein transport inhibitors, 500X) (00497593, Invitrogen) for 5 hours. In the co-culture of PBMCs with primary tumor cells or KURAMOCHI, T cell expansion media was mixed with DMEM/Ham’s F-12 (1:1). Ovaries from human female cadavers were extracted post-mortem at the Dakota Lions and Health Tissue Donation organization and stored in MACS® tissue storage solution until processed for decellularization.

### Bioengineering nECMPhys and tECMHyp 3D models

Tunable patient-derived 3D models were bioengineered by crosslinking of human plasma fibrinogen into fibrin^29, 30^. Specifically, human plasma isolated from healthy female donors or HGSC patients was mixed with Cultrex® collagen I (R&D Systems, USA) (final concentration 2 mg/ml) and trans-4- (aminomethyl)cyclohexanecarboxylic acid (T-AMCHA) as a stabilizer, and CaCl_2_ as a crosslinker in a final ratio 4:4:1:1, respectively. Unless otherwise noted, HGSC cell lines (2x10^5^ cells/ml) or primary HGSC cells (10-25 mg dissociated tissue/scaffold) were cultured inside of the 3D scaffolds in monoculture or co-culture with CAFs (2x10^5^ cells/ml). PBMCs (4x10^5^ cells/ml) were either added on top of the scaffolds and challenged to infiltrate into the matrix or multi-cultured inside the 3D scaffolds. Media was added on top of the scaffold to avoid drying and penetrated into the matrix to allow cell growth. The biomechanical properties of the model such as height, stiffness, crosslinking and stabilizing agents were optimized to recapitulate physiologically relevant oxygen levels. The 3D scaffold was incubated either in 1.5% O_2_ hypoxic incubator to recreate the oxygen levels in the hypoxic HGSC-TME originating tumor ECM Hypoxia model (tECMHyp) or in 21% O_2_ incubator recreating the physoxic oxygen levels in the normal ovary and generating the normal ECM Physoxia model (nECMPhys). When needed, the cells were isolated for further analysis by digesting the 3D scaffolds with 20 mg/ml collagenase I (17100017, Gibco) for two hours at 37°C.

### Single-cell RNA sequencing analysis

Tumor biopsy from a patient diagnosed with HGSC stage III was dissociated and primary cells were grown in co-culture with matching PBMCs in nECMPhys or tECMHyp models made with autologous plasma for 5 days. PBMCs from the same patient, not grown in the 3D scaffold, were used as a control. The 3D scaffolds were digested, and the cells were resuspended in PBS with 0.04% (w/V) bovine serum albumin. 50K cells with cell viability higher than 97% were collected to perform scRNA-seq library preparation.

Chromium Next GEM Single Cell 3’ Kit v3.1, 16 rxns (PN1000268, 10x Genomics) was used to prepare the libraries loading 16,500 cells/lane. Paired-end scRNA-sequencing was performed using Illumina NovaSeq6000. The scRNA-seq reads were aligned using CellRanger (10x Genomics, Cell Ranger, v6.0.1)^88^. Initially, CellRanger count was executed on each sample individually, followed by CellRanger aggregate to generate a consolidated expression matrix, a gene vector, and a barcode vector. The resulting expression matrix, along with the gene and barcode vectors, was imported into R (version 4.4.0) for downstream analysis using the Seurat package v. 5.0.2^89^. Quality was evaluated by the percentage of mitochondrial reads, gene counts, and number of reads. Low-quality cells with fewer than 200 and more than 8,500 unique feature counts, mitochondrial gene content exceeding 27%, or total RNA counts exceeding 89,500 were filtered out. The remaining high-quality cells were normalized using the SCTransform function (vst.flavor = “v2”) with cell cycle score regressed out.

Principal component analysis was performed for linear dimensional reduction, and the first fifty principal components were selected for downstream analysis. Uniform Manifold Approximation and Projection (UMAP) was used for non-linear dimensional reduction and visualization^90^. Graph-based clustering was performed on the UMAP embeddings to identify distinct cell populations. For cluster identification and annotation, the Monocle3 package (v. 1.3.7)^91^ was employed, and the Seurat object was converted using SeuratWrappers (v. 0.3.2). Single-cell clusters were identified and annotated based on a combination of canonical genes, cluster markers using the FindMarkers function, and unbiased cell type recognition using SingleR (v. 2.6.0)^92^. Three specific marker genes were used to confirm the clustering of each cell population using epithelial markers CLDN3, EPCAM, and KRT for HGSC; CALD1, STAR and TAGLN for CAFs; CD8A, CD8B and GZMK for CD8+ T cells; CD4, LTB and SELL for CD4+ T cells; GNLY, PRF1 and NKG7 for NK cells; CD79A, IGLC3 and MS4A1 for B cells; and CD14, CD68 and LYZ for Mφ^63, 93^. Dot plots were generated using the DotPlot function in Seurat to visualize the expression of selected marker genes across identified clusters, specifically classical hypoxic genes and hypoxic signature correlated with bad prognosis in ovarian cancer patients^31^, matrisome genes^94, 95^, CAFs genes^94, 96^, TGF-β signaling genes^97^, and activation, exhaustion and cytotoxicity genes of immune cells^63, 98^.

To identify differentially expressed genes (DEGs) in tECMHyp compared to nECMPhys within each cell type, the FindMarkers function in Seurat was used with default parameters. DEGs were defined as genes with an absolute log2-fold change greater than 0.4 and a p-value threshold lower than 0.05. Previously mentioned genes correlated with matrisome, TGF-β signaling, and activation, exhaustion, and cytotoxicity of immune cells were color-coded highlighted in orange, blue, green, purple, and red, respectively in the volcano plots and DEGs tables.

CellChat R package (v.2.1.2)^99^ was used to quantitatively infer intercellular communication networks from scRNA-seq data. Circle plots show putative ligand–receptor interactions between cells, where the thickness of the line represents the strength of the communication. Heatmaps represent the summary of the signaling pathways that contribute to outgoing or incoming communication and the color bar represents the relative signaling strength of a signaling pathway across cell types, and the bars indicate the sum of the signaling strength of each cell type or pathway. Bubble plots show the probability of interactions between receptors and ligands among different cell types with senders on y-axis and receivers on the x-axis. Weighted-directed network in CellChat was used to identify the signaling roles of the cell types represented by a heatmap that shows the senders (cell type sending signals), receivers (cell type receiving signals), mediators (cell type that controls the communication flow between the cells), and influencers (cell type that influences the information flow between cells) of intercellular communication.

### Characterization of physiologically relevant oxygen levels by optical sensor

KURAMOCHI, OVSAHO, OVCAR-3, OVCAR-8, CAOV3, OV81.2 and HEYA8 cells were multi-cultured with CAFs and PBMCs in nECMPhys and tECMHyp. The partial pressure of oxygen (pO_2,_ kPa) was measured in the middle of the 3D scaffolds on days 0, 2, 3, and 7, using the optical sensor PreSens Needle-Type Oxygen Microsensor NTH-PSt7 (Regensburg, Germany) and manual micromanipulator following manufacturer’s instructions.

### Immune infiltration assay

For immune infiltration studies, KURAMOCHI cells were monocultured or co-cultured with CAFs in nECMPhys or tECMHyp for 3 days to allow ECM remodeling and study its influence in immune infiltration. Alternatively, the 3D scaffolds were pre-treated with galunisertib (LY2157299, 10μM), a targeting drug for TGF-β signaling, to prevent collagen I expression and ECM remodeling. After three days, PBMCs were added on top of the 3D scaffold and incubated for 4 days to allow immune infiltration inside the 3D scaffolds. Additionally, dissociated tumor tissue from HGSC patients was co-cultured with CAFs in 3D model engineered with autologous HGSC plasma and the infiltration studies were performed with matching PBMCs in the same settings to recreate a precision-based preclinical model.

### Decellularization and re-cellularization of human ovaries

Whole human ovaries were decellularized by submersion in 0.1% (w/V) sodium dodecyl sulfate (SDS) in sterile PBS 1X solution with continuous rotation and daily replacement of the solution for 3 days. Ovaries were transferred to 0.05% (V/V) Triton X-100 solution in sterile PBS 1X for 3 hours, followed by several washes with sterile water for 24 hours. The tissue before and after decellularization was characterized by IHC. The decellularized matrix (dECM) was frozen in Optimal Cutting Temperature (OCT) and cryosectioned in 50 μm thick sections and placed into 8-well confocal trays. For re-cellularization, the dECM was submerged in DMEM/HAM’s F-12K media for 3 hours. Then, the media was removed and KURAMOCHI cells and CAFs were added on top of dECM in 20 µl for 5 hours to allow attachment of the cells to the matrix. More media was added on top and cultured for 3 days in ndECMPhys (normal decellularized ECM in physoxia) or in tdECMHyp (tumor decellularized ECM in hypoxia) in the presence or absence of galunisertib (10μM). On day 3, PBMCs were added on top of dECM for 4 days and immune infiltration was analyzed by confocal microscopy.

### Confocal imaging

Fluorescent images were acquired using Nikon A1R Ti2E confocal microscope with 10x magnification, 1024x1024 resolution and performing z-stacks with 0.4 μm step size. The images were processed with NIS-Elements AR Analysis 5.21.03 Software. Hypoxic status of the cells grown in nECMPhys or tECMHyp was detected by using Image-iT^TM^ Green Hypoxia Reagent (5 μM, I14834, Invitrogen) which fluoresces in green in environments with oxygen lower than 5%. KURAMOCHI, CAFs, or PBMCs in monoculture or their combinations in multi-culture KURAMOCHI:CAFs, or KURAMOCHI:CAFs:PBMCs were cultured in nECMPhys or tECMHyp for 7 days and the hypoxia-sensitive dye was added 24 h prior to imaging. To explore cell viability, the cells were stained with NUCLEAR-ID® blue/red cell viability reagent (ENZ-53005, Enzo) following the manufacturer’s instructions. This reagent stains the live cells in blue (permeable nucleic acid stain) and dead cells in red (impermeable nucleic acid stain). For visualization purposes, the blue color of live cells was modified to green color. The number of live and dead cells was manually counted with Cell Counter Plugin in ImageJ and the cell death was expressed as the percentage of dead (red) cells of total number of cells. For immune infiltration studies in nECMPhys and tECMHyp, the cells were stained 24 hours prior imaging with antibodies to detect KURAMOCHI cells (MUC1+, magenta), CAFs (CD90+, red), and infiltrated PBMCs (CD45+, green) (Table 2). For immune infiltration studies in decellularized matrix from human ovaries, the cells were pre-stained with lipophilic cell surface dyes, KURAMOCHI (DiD+, magenta), CAFs (DiI+, red), and PBMCs (DiO+, green). The z-stack image of 3D view was processed to a 2D maximum projection, which takes all the 3D data and turns it into a single 2D image for better visualization and quantification purposes. To count the number of infiltrated PBMCs (green), automatic binary thresholding was used to detect the cells in green channel with area restriction lower than 50% of the maximum area detected for all objects (0-25 μm^2^) to avoid the counting of unspecific stains.

**Table 2.** List of antibodies/staining dyes and cytokines used in the study.

| Technique | Assay | Antibody | Source | Product | Dilution |
| --- | --- | --- | --- | --- | --- |
| **IHC** | 3D models | anti-HIF1α | Rabbit | HPA001275, Sigma Aldrich | 1:100 |
|  |  | anti-collagen I | Rabbit | HPA011795, Sigma Aldrich | 1:50 |
| | | anti-α-SMA | Mouse | NBP2-33006, Novus | 1:100 |
|  | dECM | anti-collagen I | Rabbit | HPA011795, Sigma Aldrich | 1:50 |
|  |  | anti-fibronectin | Rabbit | ab 2413, Abcam | 1:100 |
| **Confocal Microscopy** | Infiltration in 3D scaffolds | APC-anti-MUC-1 | Mouse | 355608, BioLegend | 5 μl/sample |
| | | BV605-anti-CD90 | Mouse | 328128, BioLegend | 5 μl/sample |
| | | FITC-anti-CD45 | Mouse | 304038, BioLegend | 5 μl/sample |
|  | Infiltration in dECM | Invitrogen™ DiD | N/A | D307, Invitrogen | 1:1000 |
|  |  | Invitrogen™ DiI | N/A | D282, Invitrogen | 1:1000 |
|  |  | Invitrogen™ DiO | N/A | D3898, Invitrogen | 1:1000 |
| **Western blot** | Canonical signaling | anti-TGF-β (56E4) | Rabbit | 3709S, Cell Signaling | 1:1000 |
|  |  | anti-SMAD 2/3 (D7G7) | Rabbit | 8685, Cell Signaling |  |
|  |  | anti-phospho-SMAD2 (Ser 465/467)/ SMAD3 (Ser423/425) | Rabbit | 8828, Cell signaling |  |
|  |  | anti-SMAD4 | Rabbit | 38454, Cell Signaling |  |
|  | Non-canonical signaling | anti-Akt (pan) (C67E7) | Rabbit | 4691S, Cell Signaling |  |
|  |  | anti-phospho-Akt (Ser473) (D9E) | Rabbit | 4060S, Cell Signaling |  |
|  |  | anti-p44/42 MAPK (Erk1/2) (137F5) | Rabbit | 4695S, Cell Signaling |  |
|  |  | anti-phospho-p44/42 MAPK (Erk1/2) (Thr202/Tyr204) (D13.14.4E) | Rabbit | 4370S, Cell Signaling |  |
| | | Anti-β-actin (13E5) | Rabbit | 4970S, Cell Signaling | |
|  |  | Anti-rabbit IgG, HRP-linked secondary | Goat | 7074, Cell Signaling | 1:5000 |
|  | Metalloproteinase-9 | Anti-MMP-9 | Rabbit | MA5-32705, ThermoScientific | 1:1000 |
| Imaging flow cytometry | TGF-β | Anti-TGF-β (3C11) | Mouse | sc130348, Santa Cruz | 5 μl/sample |
| | | CF<sup>TM</sup> 488A anti-mouse IgG (H+L) secondary antibody | Goat | SAB4600042, Sigma-Aldrich | 1 μl/sample |
| | Collagen I | AF488-anti-collagen I | Rabbit | ab275996, Abcam | 5 μl/sample |
| | Both assays | APC-anti-FAP | Mouse | FAB3715A, R&D Systems | 5 μl/sample |
| Flow cytometry | HIF-1a | AF647-anti-HIF-1α antibody | Mouse | 359706, BioLegend | 5 μl/sample |
| | Immune Functional Assays | AF700-anti-CD3 | Mouse | 300424, BioLegend | 1 μl/sample |
| | | FITC-anti-CD4 | Mouse | 300538, BioLegend | 5 μl/sample |
| | | Perc/Cy5.5-anti-CD8 | Mouse | 344710, BioLegend | 5 μl/sample |
| | | AF647-anti-FOXP3 | Mouse | 320114, BioLegend | 1 μl/sample |
|  |  | APC-anti-CD69 | Mouse | 310910, BioLegend |  |
|  |  | BV605-anti-CD25 | Mouse | 356142, BioLegend |  |
| | | BV421-anti-IFNγ | Mouse | 502532, BioLegend | |
|  |  | PE/Dazzle<sup>TM</sup> 594 anti-IL-2 | Rat | 500344, BioLegend |  |
|  |  | BV421-anti-CD223 (LAG3) | Mouse | 369314, BioLegend |  |
|  |  | PE/Dazzle 594<sup>TM</sup>-anti-PD-1 | Mouse | 329940, BioLegend |  |
|  |  | PE-Cy7-anti-granzyme B | Mouse | 372214, BioLegend |  |
|  | Immunophenotyping | BV421-anti-CD276 (B7-H3) | Mouse | 565829, BD Horizon |  |
|  |  | PE-anti-FAP | Mouse | FAB3715P, R&D Systems |  |
|  |  | BV711-anti-CD3 | Mouse | 300464, BioLegend |  |
|  |  | PE-Cy5-anti-CD4 | Mouse | 317412, BioLegend |  |
|  |  | APC-Cy7-anti-CD8 | Mouse | 344714, BioLegend |  |
|  |  | BV785-anti-CD14 | Mouse | 301840, BioLegend |  |
|  |  | APC-anti-CD19 | Mouse | 302212, BioLegend |  |
|  |  | PE-Cy7-anti-CD56 | Mouse | 318318, BioLegend |  |
|  |  | FITC-anti-CD11b | Mouse | 301330, BioLegend |  |
| | Apoptosis in 3D cultures | FITC Annexin V Apoptosis detection kit with PI | N/A | 640914, BioLegend | 5 μl/sample |
| | | BV510-anti-CD45 | Mouse | 304036, BioLegend | 1 μl/sample |
|  | Cytotoxic effect in 2D cultures | Invitrogen™ DiO | N/A | D3898, Invitrogen | 1:1000 |
| | | PerCP/Cy5.5-anti-CD8 | Mouse | 344710, BioLegend | 1 μl/sample |
| | | PE-Cy7-anti-granzyme B | Mouse | 372214, BioLegend | 1 μl/sample |
| **Procartaplex Cytokine Array** | Pro-tumoral | IL-1β, IL-6, IL-8, macrophage inflammatory protein 1β (MIP-1β), granulocyte colony stimulating factor (G-CSF), granulocyte-macrophage colony stimulating factor (GM-CSF), vascular endothelial growth factor A (VEGF-A) | N/A | PPX45MXGZF4J, Thermo Scientific | N/A |
| | Anti-tumoral | IL-2, IL-7, IL-17A, interferon $\gamma$ (IFNγ), tumor necrosis factor alpha (TNFα) | N/A | PPX45MXGZF4J, Thermo Scientific | N/A |

### Immunohistochemistry (IHC)

For immunohistochemistry, the KURAMOCHI cells, CAFs, PBMCs or their co-cultures were seeded inside of the 3D scaffolds in BEEM® Embedding Capsules (1001, Spi Supplies). For re-cellularized dECM assessment, KURAMOCHI cells and CAFs were co-cultured on top of the dECM for 3 days. Then, the 3D scaffolds and dECM were fixed in 10% neutral buffered formalin, paraffin embedded and processed on a Leica 300 ASP tissue processor. The 3D models were oriented with the top facing forward to identify the top and the bottom of the model, and the blocks were serially sectioned at 10 µm. The sections were stained with the Discovery Ultra automated slide staining system (Roche Tissue Diagnostics) with specific antibodies (Table 2). The nECMPhys and tECMHyp scaffolds were stained for HIF-1α (master transcription regulator in response to hypoxia), extracellular and intracellular collagen I, or α-smooth muscle actin antibody (α-SMA) as a marker for CAF activation. The native ovarian tissues and dECM were stained for collagen I and fibronectin and the re-cellularized dECM for collagen I. The antigen retrieval step was performed using the cell conditioning CC1 solution (basic pH Tris based buffer, 950124, Ventana). The OmniMap HRP detection kit (7604311, Ventana) was used with DAB as the chromogen and the slides were counterstained with hematoxylin. Omission of the primary antibody served as a negative control. Moreover, native ovarian tissues and dECM were stained with Masson’s trichrome, and hematoxylin and eosin (H&E) following standard protocols. The slides were imaged in brightfield on Aperio VERSA Scanning system (Leica BioSystems) with 20x magnification and analyzed with Aperio ImageScope x64 software, detecting the signal intensity using the positive pixel count v9 algorithm in at least five randomly picked regions of interest (ROI) per section.

### Second harmonic generation microscopy (SHG)

KURAMOCHI cells or CAFs were monocultured or co-cultured in nECMPhys or tECMHyp for 7 days. Second harmonic generation microscopy (SHG) was performed to detect collagen fibers with FLUOVIEW FVMPE-RS Multi Photon Laser Scanning Microscope (Olympus) with a tunable laser (InSight, SpectraPhysics) at 840 nm. The samples were imaged using 25x water-immersion objective in two different positions of the 3D model and acquiring z-stacks of 50 μm with a step size of 1 μm. Scan size was set to 800x800 px with line averaging of 2. Six iterations of constrained iterative deconvolution were uniformly applied to the images in CellSens analysis software (Olympus). OrientationJ for Hue, Saturation and Brightness (HSB) color-survey maps and OrientationJ distribution measurement plugins in ImageJ (FIJI) were used to determine the orientation of the collagen fibers.

### Atomic force microscopy mechanical characterization

KURAMOCHI cells were grown in co-culture with CAFs in nECMPhys or tECMHyp models for 7 days. A BioScope Resolve atomic force microscopy (AFM) (Bruker) in contact mode and DNP cantilever D (Bruker, nominal spring constant: 0.06 N/m) was used to collect force-indentation curves. The actual spring contact of each cantilever was measured on a flat, hard surface using the thermal noise method. On each sample, a force-volume mode was used to collect 4x4 force-indentation curves over areas of 2x2 µm^2^. All samples were measured in PBS buffer to minimize capillary force in air. All force curves were recorded by applying a loading force of 3 nN, with a constant retraction speed of 1 µm/s, a ramp size of 5 µm, and 0.1 s of surface delay. The force-indentation curves were analyzed using the NanoScope analysis software (v1.9, Bruker). The retract part of the curve was fitted with the Hertz model:

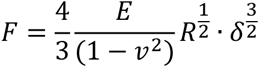

where E is the Young’s modulus, δ is the indentation depth, v is the Poisson ratio (0.3 was used here), and R is the radius of the tip curvature (20 nm).

### Imaging flow cytometry

Imaging flow cytometry was applied to quantitatively determine the intracellular TGF-β and collagen I expression. KURAMOCHI, CAFs or their co-culture were maintained for 7 days in nECMPhys and tECMHyp in the presence or absence of galunisertib (10 μM), to further explore its mechanistic properties and influence in ECM remodeling through collagen synthesis. On day 7, the 3D scaffolds were digested, and the cells were isolated, fixed and permeabilized with Cyto-Fast^TM^ Fix/Perm Buffer Kit (426803, BioLegend), and stained with antibodies (Table 2) to measure intracellular TGF-β or collagen I (green) and to distinguish CAFs (FAP+, red) from KURAMOCHI cells (FAP-) in the co-culture. Samples were analyzed at low speed and 40x magnification with Cytek® Amnis® ImageStream®X MkII Imaging Flow Cytometer (Cytek Biosciences) and INSPIRE software. Images were acquired with a 488 nm laser power of 20 mW (TGF-β) or 50 mW (collagen I), a 642 nm laser at 125 mW (collagen I) or 150 mW (TGF-β), and a 785 nm laser at 0.25 mW and in brightfield. The images were analyzed using IDEAS v6.2 analysis software. Each sample was processed by first using the area of the brightfield and the brightfield aspect ratio to identify single round cells discarding debris and speed beads. Then, the brightfield gradient RMS value was used to gate in focus events for further analysis and the area versus FAP intensity was used to distinguish CAFs from KURAMOCHI cells. Finally, events with no Mean Pixel FITC intensity were excluded from the analysis discarding debris. The area of each cell was calculated using a brightfield-based object mask and the intensity of collagen I or TGF-β was calculated from the pixel intensity within the cell mask. Finally, the mean fluorescence intensity (MFI) was analyzed in individual cells in FlowJo_v10.8.1_CL software and representative histograms were generated for each condition (Extended Data Fig. 7c).

### Western blot

KURAMOCHI cells, CAFs, or their co-culture were grown in the nECMPhys or tECMHyp for 7 days in the presence or absence of galunisertib to analyze its influence in TGF-β downstream signaling. To explore the role of oxygen levels and different cell types in metalloproteases expression, PBMCs, KURAMOCHI, CAFs, KURAMOCHI:CAFs and KURAMOCHI:CAFs:PBMCs were grown for 7 days in the nECMPhys and tECMHyp. On day 7, the above mentioned conditions were lysed with RIPA Buffer (R0278, Sigma-Aldrich) with addition of phenylmethulsulfonyl fluoride 1 mM (PMSF, 36978, Thermo Scientific), dithiotreitol 0.5 mM (DTT, D0632, Sigma-Aldrich) and phosphatase inhibitor cocktail 2 (P5726, Sigma-Aldrich) and phosphatase inhibitor cocktail 3 (P0044, Sigma-Aldrich). The samples were sonicated on ice and centrifuged at 16,000 g for 15 min at 4°C and the concentration of proteins was determined by Pierce 660 nm Protein Assay Kit (PI22662, Thermo Scientific) following the manufacturer’s instructions. Gel electrophoresis was used in 10 or 15% Mini-Protean TGX Precast Polyacrylamide gels (Bio-Rad) and transferred to PVDF membranes using Trans-Blot Turbo RTA Mini 0.2 µm PVDF Transfer Kit (1704272, Bio-Rad). The membranes were blocked in Pierce Clear Milk Blocking Buffer and then incubated with antibodies (Table 2) for metalloproteinase-9 (MMP-9), and canonical (TGF-β, SMAD 2/3, phospho(p)- SMAD2/SMAD3, SMAD4) and non-canonical (Akt/pAkt, MAPK/pMAPK) TGF-β signaling. β-actin was used as a loading control. Anti-rabbit IgG, HRP-linked secondary antibody and Immobilon® Western Chemiluminescent HRP Substrate were used to detect specific bands on ChemiDoc MP Imaging System (Bio-Rad). The images were processed in Image Lab Software (Bio-Rad).

### Flow Cytometry

To detect HIF-1α expression and to perform immune functional analysis in patient-derived cultures, primary HGSC cells were co-cultured with matching PBMCs for 2 days inside the nECMPhys or tECMHyp generated with autologous plasma. The PBMCs were pre-stimulated for 5 hours for immune functional characterization. On day 2, the 3D scaffolds were digested, and the isolated cells were stained with LIVE/DEAD^TM^ Fixable blue dead cell stain kit (L23105, Invitrogen) and fixed and permeabilized with Cyto-Fast^TM^ Fix/Perm Buffer Kit (426803, BioLegend). Cells were blocked with Human TruStain FcX^TM^ (422302, BioLegend) and stained with antibody for HIF-1α or with panels of antibodies to detect intracellular cytokines or activation and exhaustion surface markers (Table 2). Moreover, immunophenotyping studies were performed to identify different immune cell populations that have infiltrated inside the 3D scaffold. For that purpose, on day 7, the isolated cells were stained with LIVE/DEAD^TM^ Fixable blue dead cell staining and surface marker antibodies (Table 2) to detect cancer cells (B7H3+), CAFs (FAP+), CD3+, CD4+, CD8+ T lymphocytes, macrophages (CD14+), B cells (CD19+), NK cells (CD56+), and myeloid-derived suppressor cells (MDSC, CD11b+). Furthermore, the cytotoxic effect of PBMCs on HGSC cells was also determined by flow cytometry. KURAMOCHI cells were mono-cultured or co-cultured inside the nECMPhys and tECMHyp with stimulated PBMCs. On day 5, the 3D scaffolds were digested, and the isolated cells were stained with antibodies to detect PBMCs (CD45+) and KURAMOCHI (CD45-). The cytotoxic effect was evaluated by staining with propidium iodide (PI) and Annexin V (Table 2) to detect live, early apoptotic and necrotic cells. Alternatively, the cytotoxic effect of the immune cells on HGSC was evaluated in 2D cultures of KURAMOCHI cells stained with cell surface marker DiO (green) co-cultured with PBMCs at increasing ratios (1:1, 1:2, 1:3 and 1:4) in normoxia (21% O_2_) or hypoxia (1.5% O_2_) for 2 days. On day 2, the cells were stained with LIVE/DEAD^TM^ Fixable blue dead cell stain kit and antibodies for CD8 and Granzyme B (Table 2). Flow cytometry was performed using BD FACS Fortessa and FACSDiva v6.1.2 software by acquiring a minimum of a ten thousand Precision Count Beads^TM^ (424902, Biolegend, USA). The data was analyzed using FlowJo v10 software (Ashland, OR). HIF-1α expression was analyzed by manual gating and cell counts were normalized to counting beads. For immune functional assays and immunophenotyping studies, single live cells were analyzed by dimensionality reduction analysis using Phenograph v2.5.0 and UMAP v4.1.1. (default settings) plugins in FlowJo software. The level of expression of the markers in functional assays was determined as the frequency (percentage of the CD4+ or CD8+ parent population) and the stacked charts expressed the percentage of each subpopulation in the total population of CD4+ or CD8+ T cells, with a total of 100%. The different cluster populations of infiltrated lymphocytes were identified with different colors and numbers and expressed as counts of cells positive for selected marker inside the 3D scaffold. The cytotoxic effect of PBMCs on KURAMOCHI cells in the 3D scaffolds was expressed as a percentage of KURAMOCHI cells in each stage (live, early apoptosis and necrosis), and the cytotoxic effect in 2D was determined as the fold change of live KURAMOCHI cells in the absence of PBMCs.

### ProCartaPlex Immunoassays

Customized ProcartaPlex (PPX45MXGZF4J, Thermo Scientific) was used following the manufacturer’s instructions to detect extracellular pro-tumoral, anti-tumoral cytokines, and growth factors (Table 2) from supernatants of digested 3D scaffolds and culture media mixed in 1:1 ratio and collected from infiltration assays in patient-derived model. The samples were analyzed with Luminex 200 analyzer and Belysa® Immunoassay Curve Fitting Software using 4-PL Sigmoidal curve extrapolation to detect the concentration (pg/ml) of each analyte based on standards. The data was standardized by calculating the z-score by subtracting the mean of all samples divided by the standard deviation of all samples for each analyte. The heatmaps were generated using the median of the z-score of six patients per condition.

### ELISA

The cultures of PBMCs, KURAMOCHI, CAFs, KURAMOCHI:CAFs or KURAMOCHI:CAFS:PBMCs were grown for 7 days in the nECMPhys or tECMHyp. Extracellular metalloproteinase 9 (MMP-9) was detected in the supernatants from the digested 3D scaffolds and the culture media (mixed in 1:1) using Human MMP-9 ELISA Kit-Quantikin (DMP900, R&D Systems) following the manufacturer’s instructions. 4-PL Sigmoidal curve extrapolation was used to calculate the concentration (ng/ml) of MMP-9 based on standards.

### Statistical analysis

Experiments were performed in triplicates and repeated at least three times when cell lines were used. Unless otherwise stated, primary cells (dissociated tumors or PBMCs) included a sample size of at least six individuals per experimental group. All graphs were generated and analyzed using GraphPad Prism 9 software as paired dot plots or representing Mean ± SD, and statistical significance was analyzed using paired, unpaired t-test, or ANOVA multi-comparisons Tukey test. The data that did not follow a normal (Gaussian) distribution were graphed as a violin plot with median where statistical significance was determined with Mann-Whitney nonparametric test. p-value lower than 0.05 was considered significant.

## Data Availability

The single-cell RNA sequencing data are available in the GEO database under accession code GSE299200. All relevant data supporting the findings of this study are available within the paper and its Supplementary Information. Source data for the figures are provided with this paper.

## Code availability

Seurat and Cell Chat are open-source R packages designed for quality control, analysis and visualization of cell-cell communications from scRNA-seq data, and the tutorials and code are available in GitHub

## Acknowledgements

We gratefully acknowledge Dr. Michael Kareta and Dr. Malini Mukherjee from Functional Genomics and Bioinformatics Core in Sanford Research and Jessica Zylla from Department of Biomedical Engineering, University of South Dakota for helping with scRNA sequencing. We thank Claire Evans from Histology & Imaging Core in Sanford Research for processing and staining samples for IHC. We thank Marcy Dimond from Dakota Lions Sight and Health for providing human ovaries. This project used Sanford Research Histology and Imaging Core and Flow Cytometry Core Facilities that are supported in part by a Cancer COBRE grant from the National Institutes of Health (P30GM145398) and the Functional Genomic and Bioinformatics Core that is supported by Pediatric COBRE grant from the National Institutes of Health (P30GM154633). This work was funded by the National Cancer Institute, R21CA259158 (P.P.), R37CA291976 (P.P), and American Cancer Society, RSG-23-1149466-01-CDP (P.P). We would also like to thank the participants and volunteers, the Sanford Blood Biobank, and the clinical coordinators involved in this study. The schemes in Fig. 1a, 3a, 4a, 6a, and Extended Data Fig. 8a were generated with Biorender.com.

## Author contributions

P.P. conceived and coordinated the study. S.P. and P.P. wrote and edited the manuscript. S.P and P.P. were involved in designing, performing and analyzing experiments in the manuscript. H.A., K.C., O.I., J.W., S.B., E.Z.G., D.M.F., C.W., provided critical expertise and resources. S.P., H.A., K.C., processed samples; M.B., obtained samples from patients; O.I., analyzed scRNA-seq data; J.W., analyzed high dimensional flow cytometry data; C.W.; analyzed atomic force microscopy data; All the authors discussed and read the paper.

