## Supplemental files for "Recapitulating physiologically relevant oxygen levels and extracellular matrix remodeling in patient-derived tumor-immune tunable models reveal targeting opportunities for immunologically cold high-grade serous tumors"

**Extended Fig. 1. scRNA-seq analysis of nECMPhys vs tECMHyp.** **a** Feature plots and **b** heatmap of marker genes in clustered cell populations. **c** Clusters of cell populations split by conditions. **d** Expression of marker genes in each cell population split by conditions. **e** Feature plots of hypoxia-related genes expresion.

**Extended Fig. 2. tECMHyp reveals a transcriptional signature with involvement of ECM remodeling and immune functionality.** Feature plots with expression of **a** Matrisome, CAF-associated, and TGF-β signaling genes and **b** immune activation, exhaustion, and cytotoxicity genes identified by scRNA-seq analysis. **c** Volcano plots showing DEG analysis in tECMHyp compared to nECMPhys.

**Extended Fig. 3. CAFs are the main contributors to the collagen signaling pathway involved in tumor-stroma-immune interactions.** **a** Circle plots of weights/strength of cell-cell interactions. **b** Scatter plot with dominant senders and receivers. **c** Network centrality analysis. **d** Relative contribution of each ligand-receptor (L-R) pair. **e** Probability of L-R interactions in collagen pathway in CAFs and **f** HGSC cells with the rest of cell populations.

**Extended Fig. 4. tECMHyp mimics hypoxia-induced CAF-driven ECM remodeling and functionality of TILs.** **a** Confocal microscopy images of hypoxic cells (green). **b** HIF-1α detected by flow cytometry in primary HGSC tissue or PBMCs in monocultures and **c** by IHC in multi-culture of KURAMOCHI, CAFs and PBMCs. **d&e** IHC images and quantification of collagen I and α-SMA. MMP-9 detected by **f** western blot or **g** ELISA, Mean±SD, n=2, ANOVA. **h** SHG microscopy images in monocultures and **i** quantification of the distribution of orientation of collagen fibers for all conditions.

**Extended Fig. 5. Intratumoral hypoxia induces higher functionality and cytotoxicity of PBMCs.** **a&b** Heatmaps of functionality markers expression. **c** Apoptosis evaluation of KURAMOCHI in the presence or absence of PBMCs in tECMHyp compared to nECMPhys. **d** Cytotoxicity evaluation and granzyme B expression in PBMCs cocultured with KURAMOCHI in 2D by flow cytometry. Two-way ANOVA, $*p<0.05$, $****p<0.001$.

**Extended Fig. 6. Intratumoral hypoxia induces impaired immune infiltration.** **a** Confocal images of infiltrated CD45+ T cells (green) in 3D models with HGSC (MUC1+, magenta) and their quantifications (counts). Scale=100μm. n=14. **b** Heatmap showing the expression of surface markers used to determine the cell populations in dimensionality reduction analysis of immune cell infiltration. **c** Counts of immune cells infiltrated into 3D models with KURAMOCHI in monoculture. n=8. Paired t-test, $*p<0.05$, $***p<0.001$, ns=not significant.

**Extended Fig.7. Galunisertib decreases hypoxia-induced CAF-driven TGF-β signaling and collagen expression**. **a** Western blot images of TGF-β in monocultures. **b** Western blot of canonical and non-canonical TGF-β signaling molecules in cells cultured in the presence or absence of galunisertib (Gal) in monocultures. **c** Gating strategy and mask used to detect cells in imaging flow cytometry. **d** Representative images of TGF-β (green) and **e** collagen I (green) detected by imaging flow cytometry in CAFs (FAP+, red) or KURAMOCHI (FAP-) and quantification of green mean fluorescent intensity (MFI). Scale=10μm. **f** IHC images of collagen I and quantification of positive signal. Mean±SEM. Scale=200µm. d&e Mann-Whitney test, f ANOVA multi-comparisons Tukey test, *p<0.05, **p<0.01, ***p<0.001, ****p<0.0001, ns=not significant.

**Extended Fig. 8. Preincubation with galunisertib reverses the impaired immune infiltration in the HGSC patient-derived model. a** Schematic representation of infiltration studies in the presence or absence of galunisertib (Gal). **b** Heatmaps showing the expression of the surface markers used to detect the immune cell populations in dimensionality reduction analysis by flow cytometry in the infiltration of autologous PBMCs into matching HGSC primary tumor cocultured with CAFs.

**Extended Fig. 9. Galunisertib rescues the impaired immune infiltration in the HGSC cell line model. a** Confocal microscopy images of infiltrated CD45+ PBMCs (green) in 3D models with HGSC (MUC1+, magenta) in the presence or absence of galunisertib (Gal) and their quantifications (counts). Scale=100µm, n=8. **b** Heatmaps of markers to detect immune cells in monoculture and co-culture in dimensionality reduction analysis by flow cytometry. **c** Density plots and quantification (counts) of infiltrated immune phenotypes in KURAMOCHI monocultures with or without Gal, n=13. **d** Confocal microscopy of live (green) or dead cells (red) of infiltrated PBMCs in KURAMOCHI in monoculture or co-culture with CAFs and quantification of survival (percentage of death of total number of cells). Scale=100µm. c Paired t-test. d ANOVA, *p<0.05, **p<0.01, ****p<0.0001, ns=not significant.

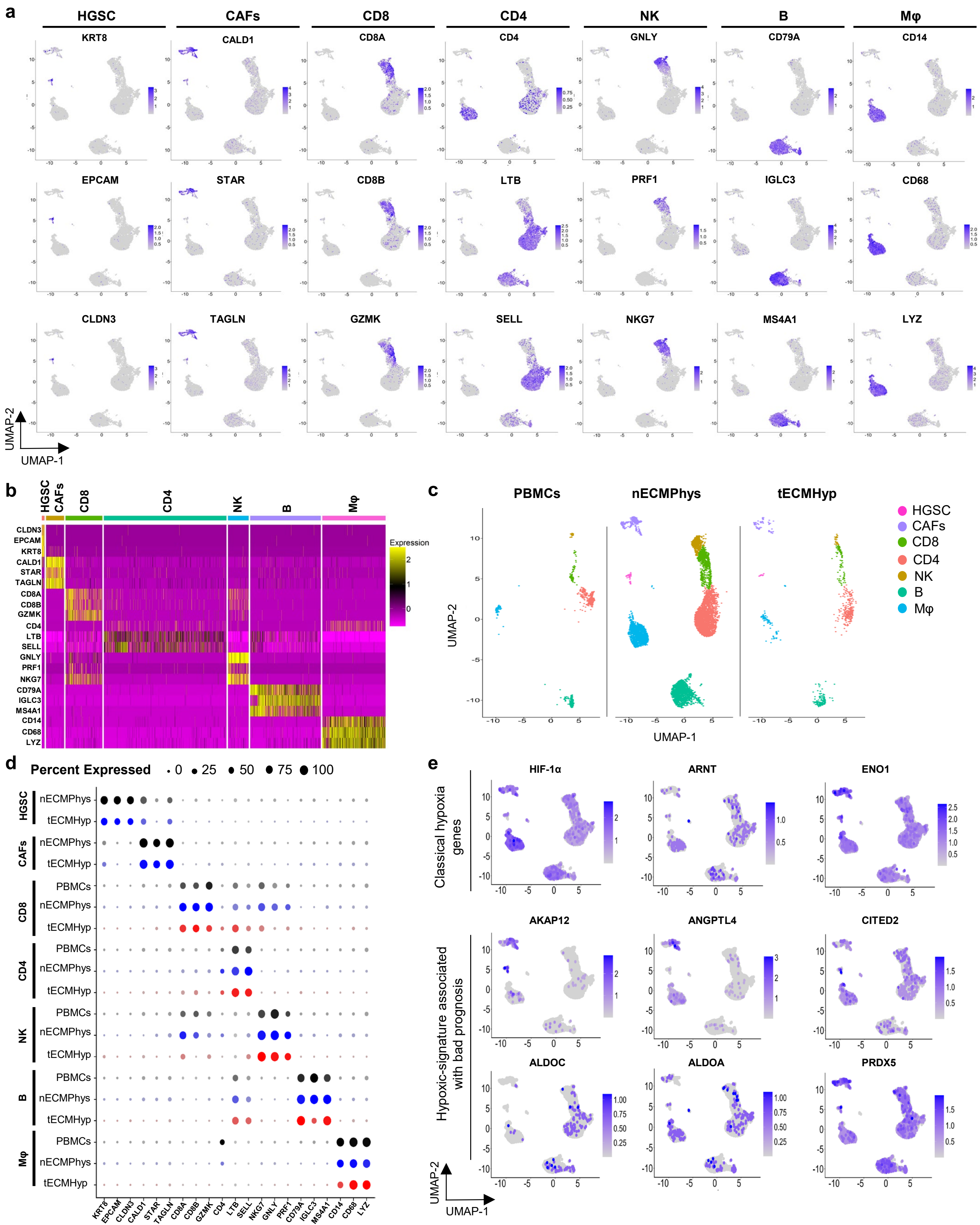

a
HGSC
CAFs
CD8
CD4
NK
B
Mφ
KRT8
CALD1
CD8A
CD4
GNLY
CD79A
CD14
EPCAM
STAR
CD8B
LTB
PRF1
IGLC3
CD68
CLDN3
TAGLN
GZMK
SELL
NKG7
MS4A1
LYZ
UMAP-2
UMAP-1
b
Expression
c
PBMCs
nECMPhys
tECMHyp
HGSC
CAFs
CD8
CD4
NK
B
Mφ
d
Percent Expressed
0
25
50
75
100
e
HIF-1α
ARNT
ENO1
Classical hypoxia genes
AKAP12
ANGPTL4
CITED2
ALDOC
ALDOA
PRDX5
Hypoxic-signature associated with bad prognosis

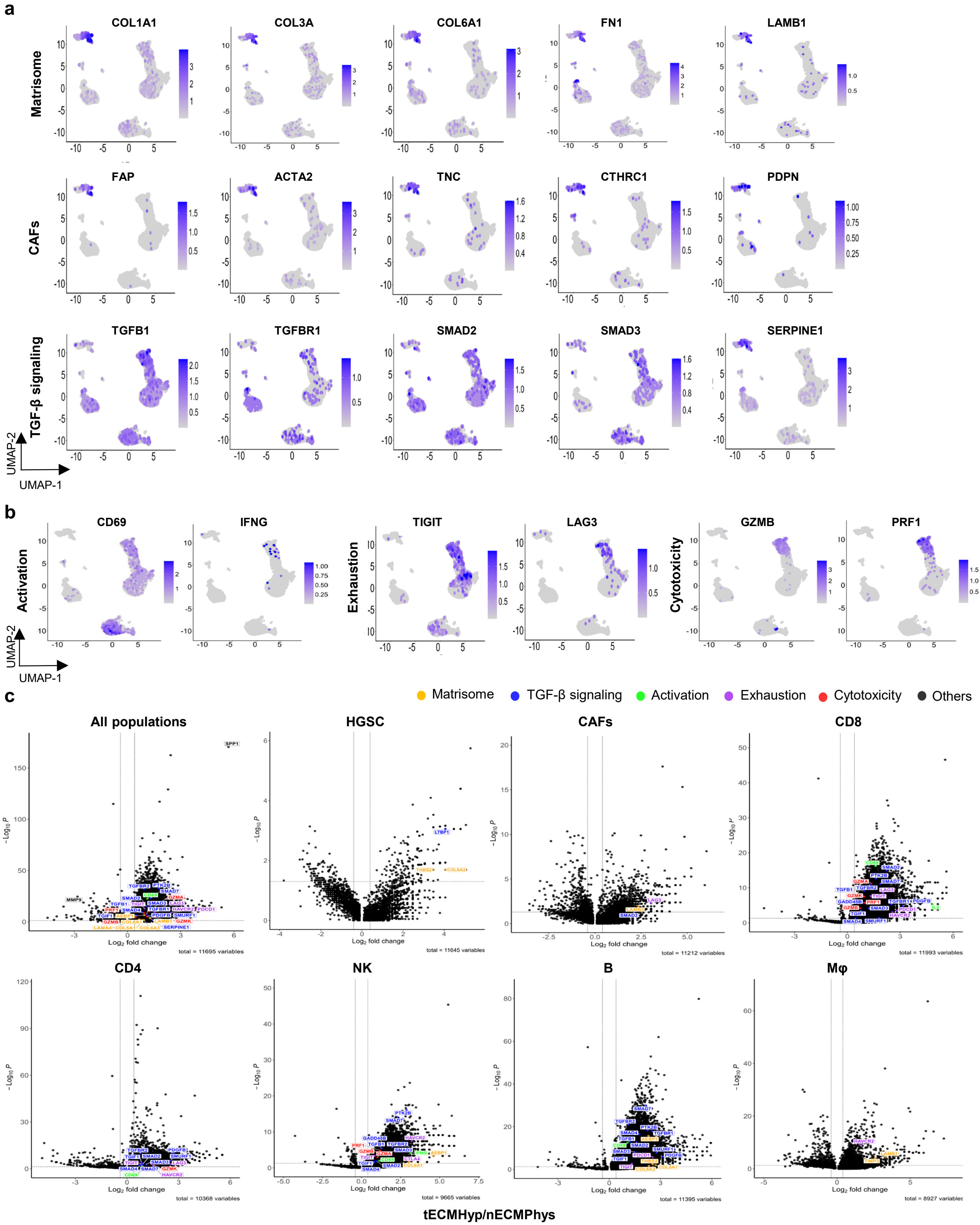

a
Matrisome
COL1A1
COL3A
COL6A1
FN1
LAMB1
CAFs
FAP
ACTA2
TNC
CTHRC1
PDPN
TGF-β signaling
TGFB1
TGFBR1
SMAD2
SMAD3
SERPINE1
UMAP-2
UMAP-1
b
Activation
CD69
IFNG
Exhaustion
TIGIT
LAG3
Cytotoxicity
GZMB
PRF1
UMAP-2
UMAP-1
c
Matrisome
TGF-β signaling
Activation
Exhaustion
Cytotoxicity
Others
All populations
HGSC
CAFs
CD8
CD4
NK
B
Mφ
Log2 fold change
-Log10 P
total = 11695 variables
total = 11645 variables
total = 11212 variables
total = 11993 variables
total = 10368 variables
total = 9665 variables
total = 11395 variables
total = 8927 variables
tECMHyp/nECMPhys

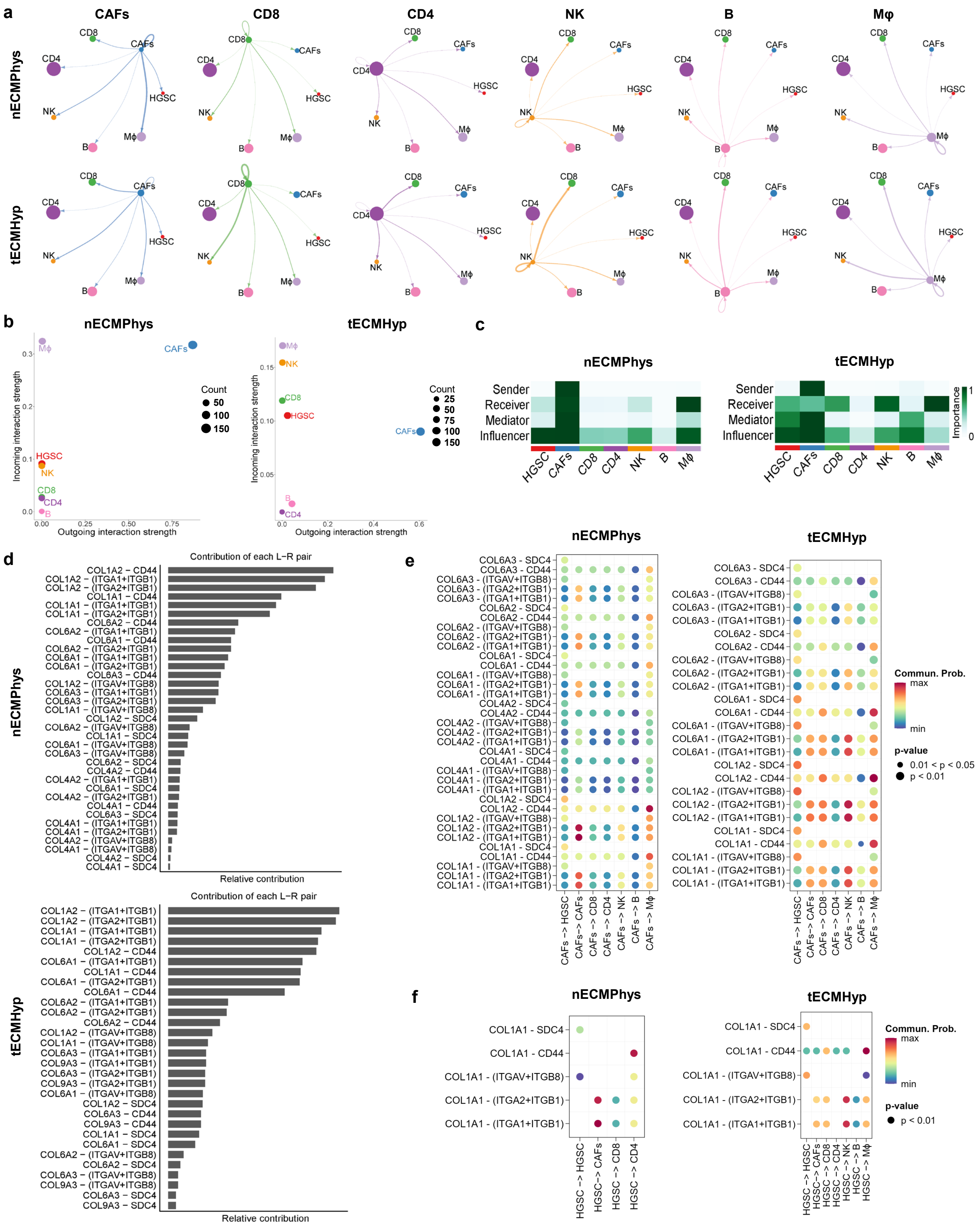

a
CAFs
CD8
CD4
NK
B
Mφ
nECMPhys
tECMHyp
b
nECMPhys
tECMHyp
Incoming interaction strength
Outgoing interaction strength
Count
c
nECMPhys
tECMHyp
Sender
Receiver
Mediator
Influencer
Importance
HGSC
CAFs
CD8
CD4
NK
B
Mφ
d
Contribution of each L-R pair
Relative contribution
e
Commun. Prob.
max
min
p-value
0.01 < p < 0.05
p < 0.01
f

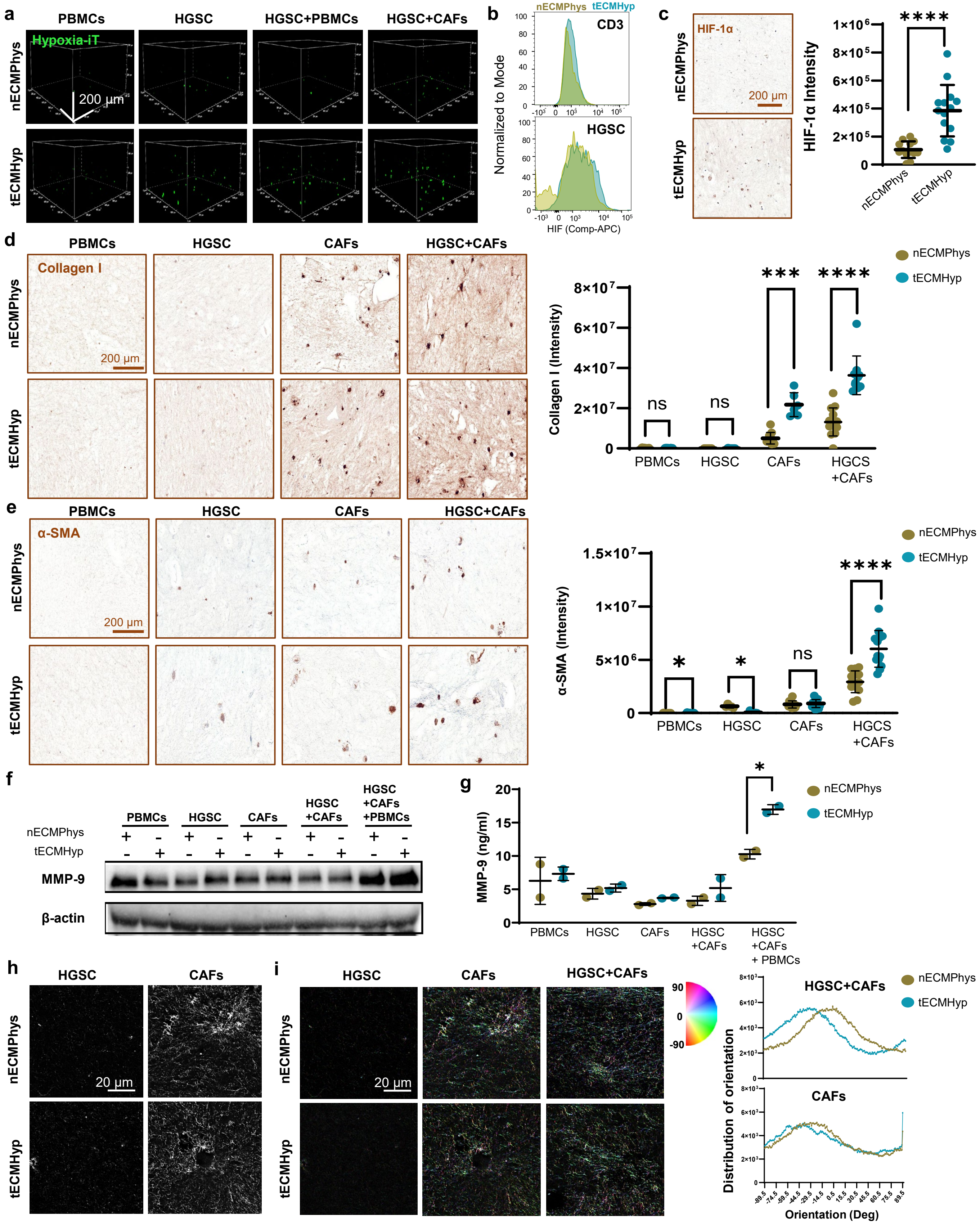

a
PBMCs
HGSC
HGSC+PBMCs
HGSC+CAFs
nECMPhys
tECMHyp
Hypoxia-iT
200 µm
b
nECMPhys tECMHyp
CD3
HGSC
Normalized to Mode
HIF (Comp-APC)
c
HIF-1α
200 µm
HIF-1α Intensity
****
nECMPhys
tECMHyp
d
PBMCs
HGSC
CAFs
HGSC+CAFs
Collagen I
200 µm
Collagen I (Intensity)
ns
ns
***
****
HGCS +CAFs
e
α-SMA
200 µm
α-SMA (Intensity)
*
*
ns
****
f
PBMCs
HGSC
CAFs
HGSC +CAFs
HGSC +CAFs +PBMCs
MMP-9
β-actin
g
MMP-9 (ng/ml)
*
HGSC +CAFs + PBMCs
h
HGSC
CAFs
20 µm
i
HGSC
CAFs
HGSC+CAFs
20 µm
HGSC+CAFs
CAFs
Distribution of orientation
Orientation (Deg)

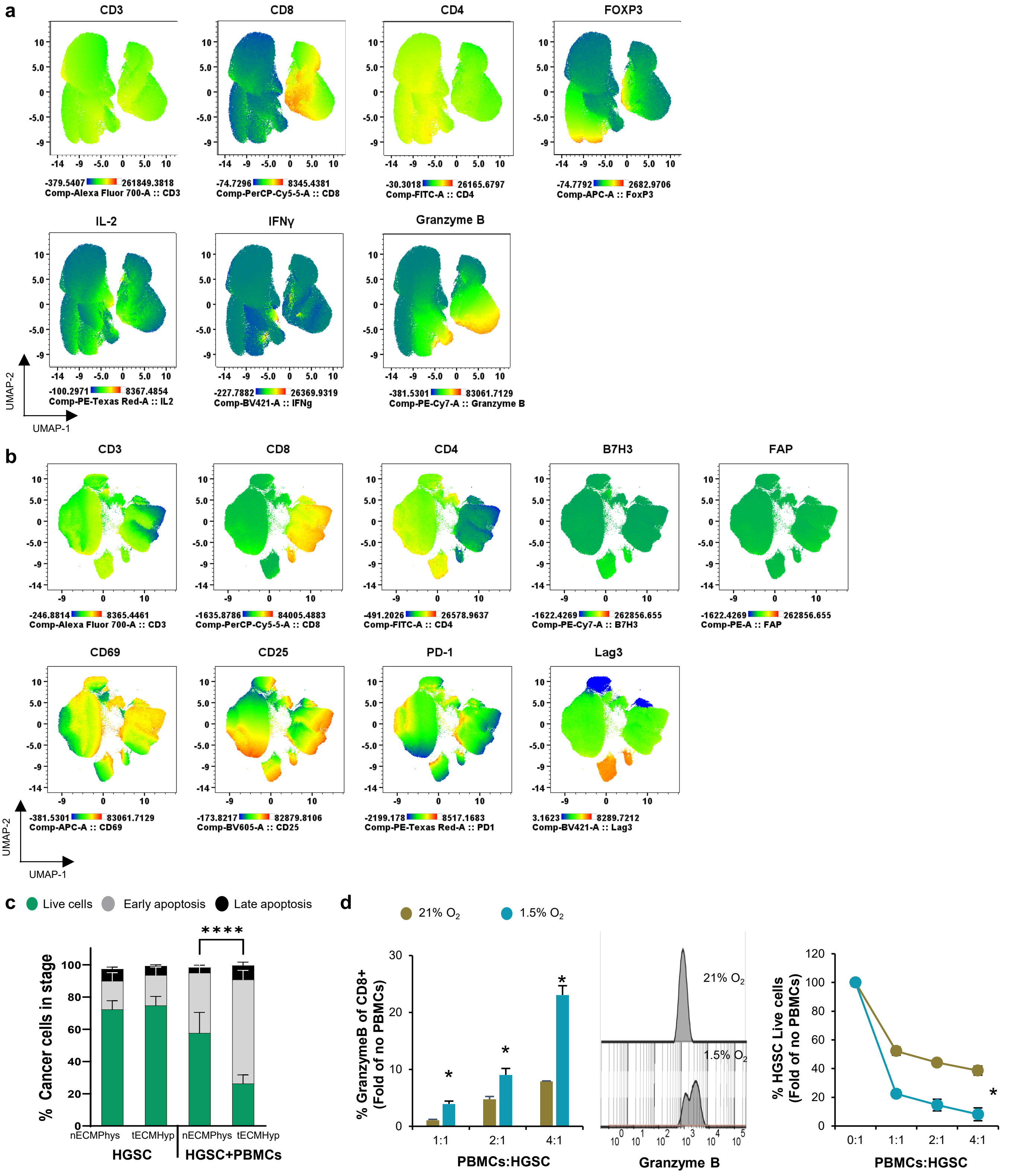

a
CD3
CD8
CD4
FOXP3
-379.5407 261849.3818
Comp-Alexa Fluor 700-A :: CD3
-74.7296 8345.4381
Comp-PerCP-Cy5-5-A :: CD8
-30.3018 26165.6797
Comp-FITC-A :: CD4
-74.7792 2682.9706
Comp-APC-A :: FoxP3
IL-2
IFNγ
Granzyme B
-100.2971 8367.4854
Comp-PE-Texas Red-A :: IL2
-227.7882 26369.9319
Comp-BV421-A :: IFNg
-381.5301 83061.7129
Comp-PE-Cy7-A :: Granzyme B
UMAP-2
UMAP-1
b
CD3
CD8
CD4
B7H3
FAP
-246.8814 8365.4461
Comp-Alexa Fluor 700-A :: CD3
-1635.8786 84005.4883
Comp-PerCP-Cy5-5-A :: CD8
-491.2026 26578.9637
Comp-FITC-A :: CD4
-1622.4269 262856.655
Comp-PE-Cy7-A :: B7H3
-1622.4269 262856.655
Comp-PE-A :: FAP
CD69
CD25
PD-1
Lag3
-381.5301 83061.7129
Comp-APC-A :: CD69
-173.8217 82879.8106
Comp-BV605-A :: CD25
-2199.178 8517.1683
Comp-PE-Texas Red-A :: PD1
3.1623 8289.7212
Comp-BV421-A :: Lag3
UMAP-2
UMAP-1
c
Live cells
Early apoptosis
Late apoptosis
****
% Cancer cells in stage
nECMPhys
tECMHyp
nECMPhys
tECMHyp
HGSC
HGSC+PBMCs
d
21% O2
1.5% O2
% GranzymeB of CD8+ (Fold of no PBMCs)
PBMCs:HGSC
21% O2
1.5% O2
Granzyme B
% HGSC Live cells (Fold of no PBMCs)
PBMCs:HGSC

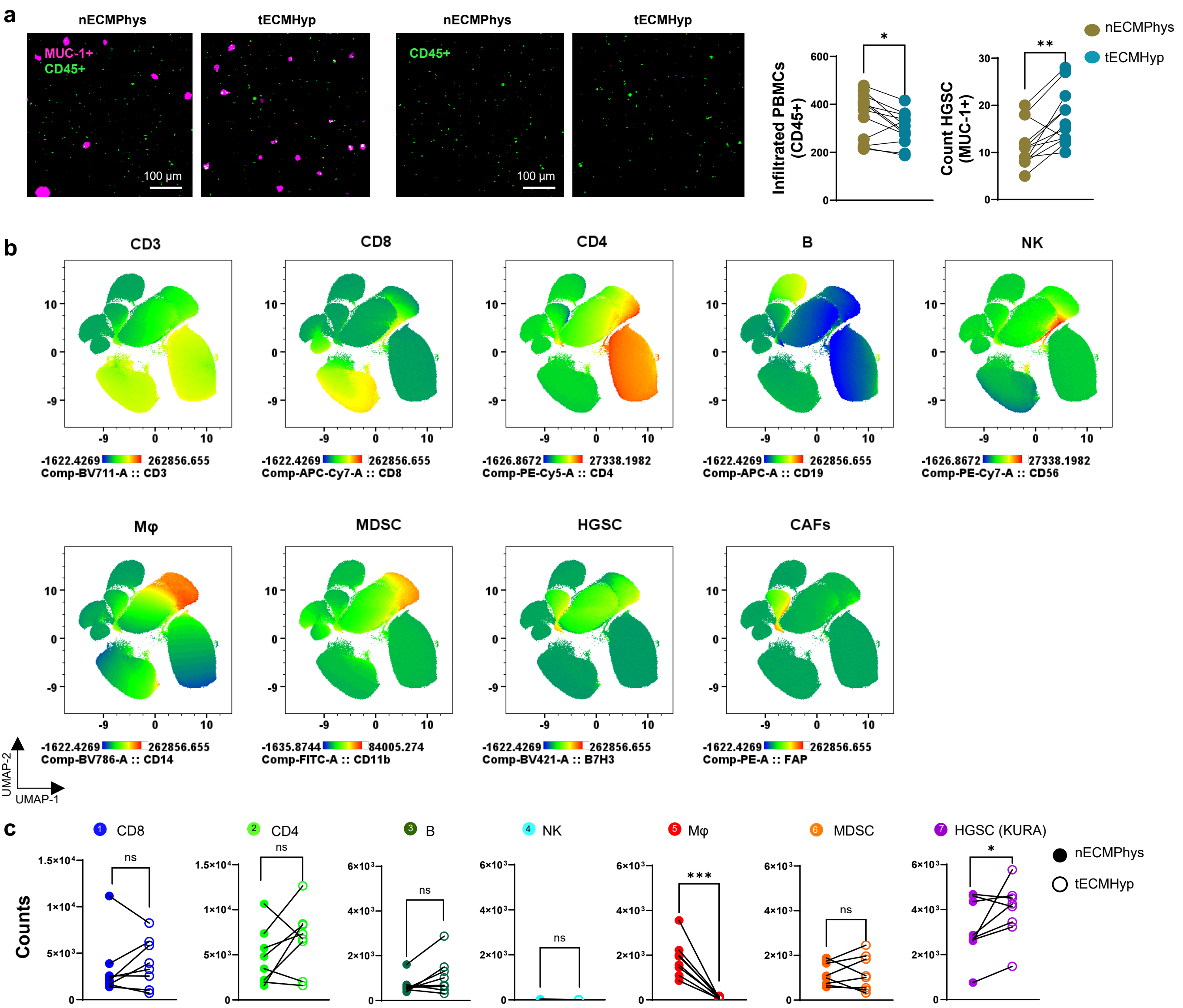
a
nECMPhys
tECMHyp
nECMPhys
tECMHyp
MUC-1+
CD45+
CD45+
100 µm
100 µm
Infiltrated PBMCs (CD45+)
Count HGSC (MUC-1+)
nECMPhys
tECMHyp
b
CD3
CD8
CD4
B
NK
-1622.4269 262856.655 Comp-BV711-A :: CD3
-1622.4269 262856.655 Comp-APC-Cy7-A :: CD8
-1626.8672 27338.1982 Comp-PE-Cy5-A :: CD4
-1622.4269 262856.655 Comp-APC-A :: CD19
-1626.8672 27338.1982 Comp-PE-Cy7-A :: CD56
Mφ
MDSC
HGSC
CAFs
-1622.4269 262856.655 Comp-BV786-A :: CD14
-1635.8744 84005.274 Comp-FITC-A :: CD11b
-1622.4269 262856.655 Comp-BV421-A :: B7H3
-1622.4269 262856.655 Comp-PE-A :: FAP
UMAP-2
UMAP-1
c
1 CD8
2 CD4
3 B
4 NK
5 Mφ
6 MDSC
7 HGSC (KURA)
Counts
nECMPhys
tECMHyp

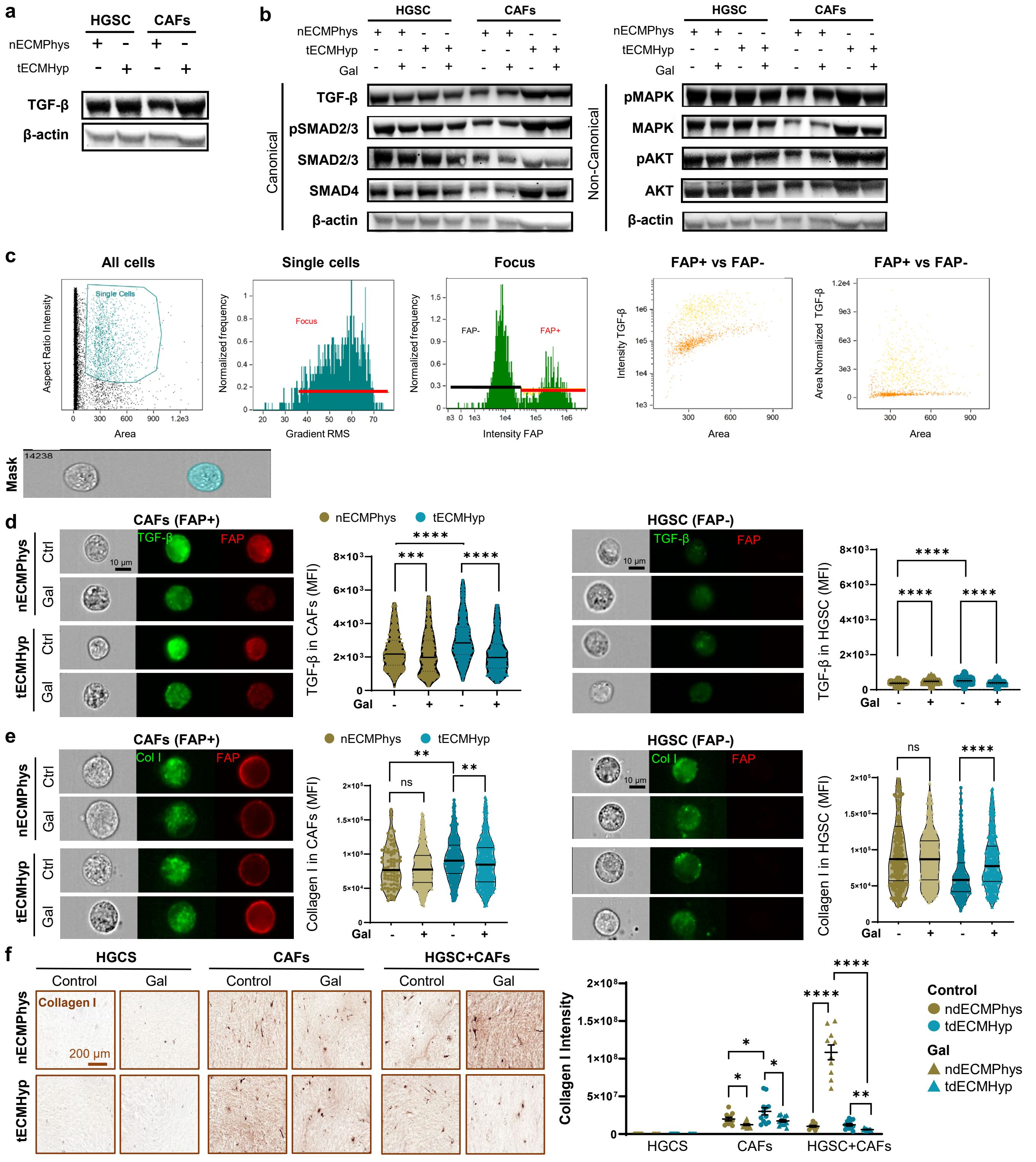

a
HGSC
CAFs
nECMPhys
tECMHyp
TGF-β
β-actin
b
HGSC
CAFs
nECMPhys
tECMHyp
Gal
Canonical
TGF-β
pSMAD2/3
SMAD2/3
SMAD4
β-actin
Non-Canonical
pMAPK
MAPK
pAKT
AKT
β-actin
c
All cells
Single Cells
Aspect Ratio Intensity
Area
Single cells
Focus
Normalized frequency
Gradient RMS
FAP-
FAP+
Intensity FAP
FAP+ vs FAP-
Intensity TGF-β
Area Normalized TGF-β
Mask
14238
d
CAFs (FAP+)
TGF-β
FAP
10 μm
nECMPhys
tECMHyp
Ctrl
Gal
TGF-β in CAFs (MFI)
HGSC (FAP-)
TGF-β in HGSC (MFI)
e
CAFs (FAP+)
Col I
Collagen I in CAFs (MFI)
HGSC (FAP-)
Collagen I in HGSC (MFI)
ns
f
HGCS
CAFs
HGSC+CAFs
Control
Gal
Collagen I
200 μm
Collagen I Intensity
ndECMPhys
tdECMHyp

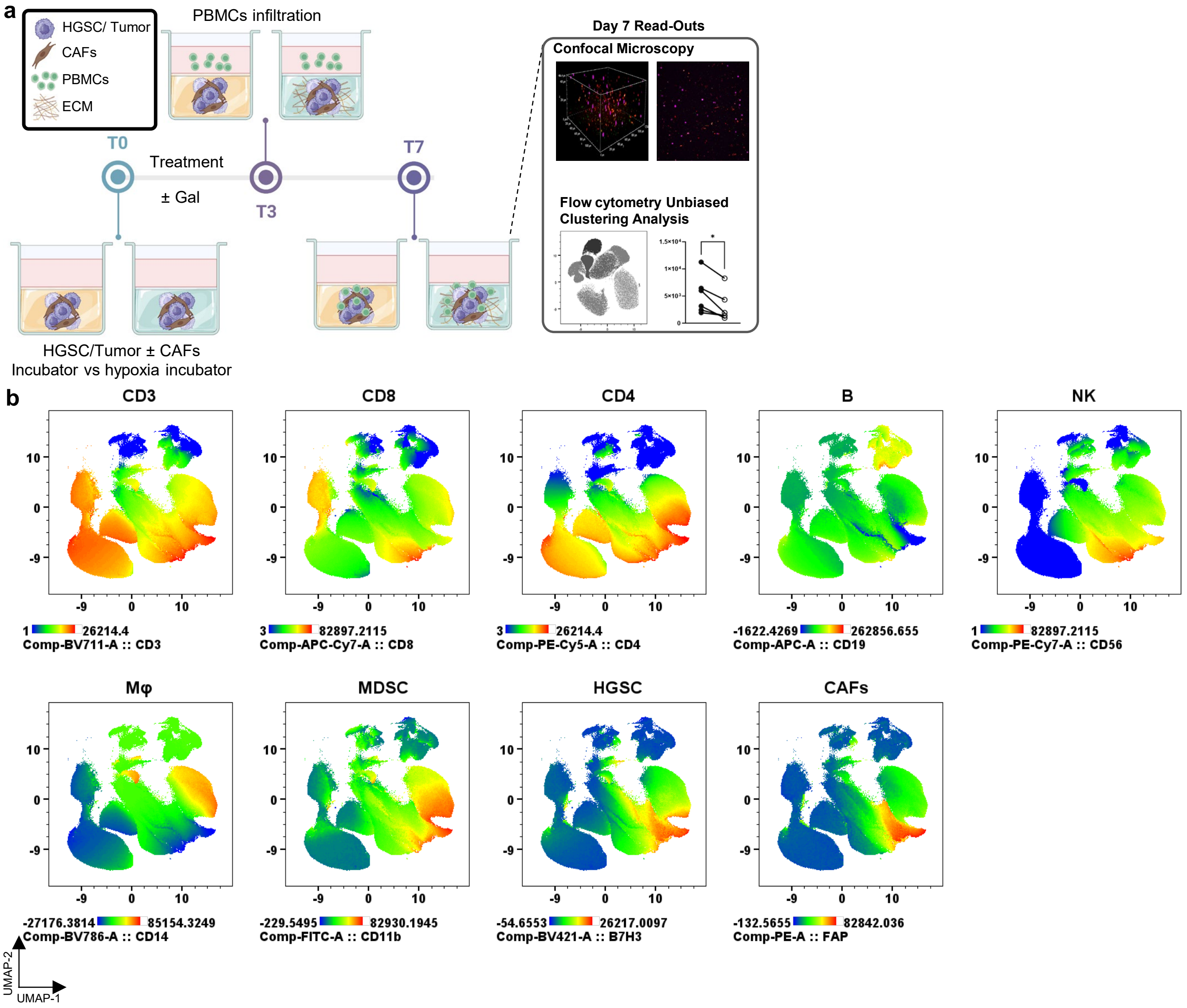
a
HGSC/ Tumor
CAFs
PBMCs
ECM
PBMCs infiltration
T0
Treatment
± Gal
T3
T7
HGSC/Tumor ± CAFs
Incubator vs hypoxia incubator
Day 7 Read-Outs
Confocal Microscopy
Flow cytometry Unbiased
Clustering Analysis
b
CD3
CD8
CD4
B
NK
1 26214.4
Comp-BV711-A :: CD3
3 82897.2115
Comp-APC-Cy7-A :: CD8
3 26214.4
Comp-PE-Cy5-A :: CD4
-1622.4269 262856.655
Comp-APC-A :: CD19
1 82897.2115
Comp-PE-Cy7-A :: CD56
Mφ
MDSC
HGSC
CAFs
-27176.3814 85154.3249
Comp-BV786-A :: CD14
-229.5495 82930.1945
Comp-FITC-A :: CD11b
-54.6553 26217.0097
Comp-BV421-A :: B7H3
-132.5655 82842.036
Comp-PE-A :: FAP
UMAP-2
UMAP-1

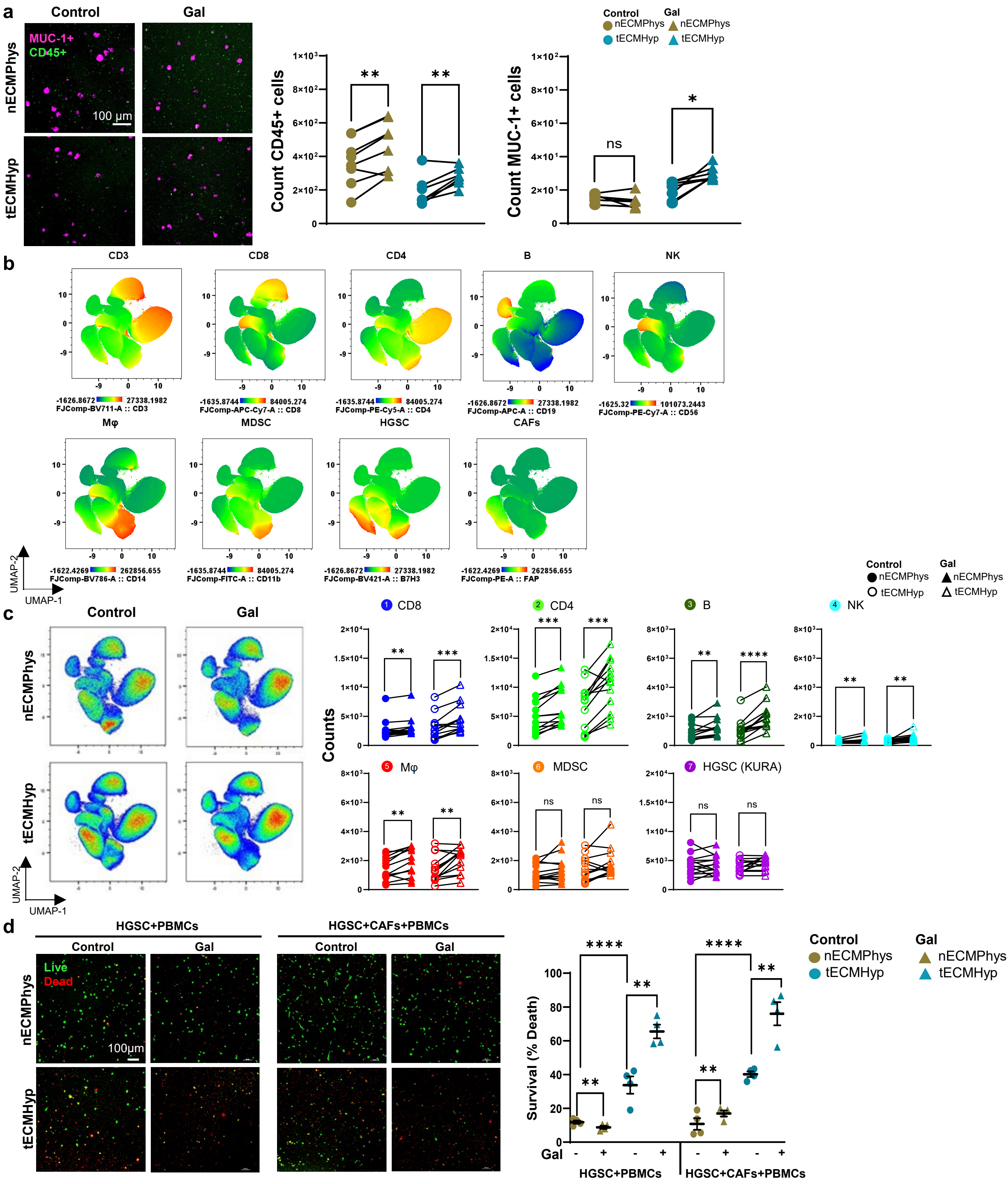

a
Control
Gal
nECMPhys
tECMHyp
MUC-1+
CD45+
100 µm
Control
Gal
nECMPhys
tECMHyp
Count CD45+ cells
Count MUC-1+ cells
ns
b
CD3
CD8
CD4
B
NK
-1626.8672 27338.1982
FJComp-BV711-A :: CD3
-1635.8744 84005.274
FJComp-APC-Cy7-A :: CD8
-1635.8744 84005.274
FJComp-PE-Cy5-A :: CD4
-1626.8672 27338.1982
FJComp-APC-A :: CD19
-1625.32 101073.2443
FJComp-PE-Cy7-A :: CD56
Mφ
MDSC
HGSC
CAFs
-1622.4269 262856.655
FJComp-BV786-A :: CD14
-1635.8744 84005.274
FJComp-FITC-A :: CD11b
-1626.8672 27338.1982
FJComp-BV421-A :: B7H3
-1622.4269 262856.655
FJComp-PE-A :: FAP
UMAP-2
UMAP-1
c
Control
Gal
nECMPhys
tECMHyp
CD8
CD4
B
NK
Counts
Mφ
MDSC
HGSC (KURA)
ns
d
HGSC+PBMCs
HGSC+CAFs+PBMCs
Control
Gal
Live
Dead
nECMPhys
tECMHyp
100µm
Survival (% Death)
Gal
HGSC+PBMCs
HGSC+CAFs+PBMCs
Control
Gal
nECMPhys
tECMHyp
